# Dynamic environments require photosynthetic electron flows with distinct bandwidths

**DOI:** 10.1101/2025.05.16.654609

**Authors:** Sai Kiran Madireddi, Liat Adler, Carolyne Stoffel, Myria Schröder, Dimitri Tolleter, Adrien Burlacot

## Abstract

All living cells depend on the dynamic balance between their energy supply and demand to survive and thrive in dynamic environments. In extreme cases, like for photosynthetic organisms, their energy source, light, can fluctuate dramatically in intensity over timescales of seconds to hours. While various photosynthetic electron flows (EF) are crucial for maintaining bioenergetic homeostasis, how EFs are modulated to respond to dynamic energy intake remains unclear. Here, we show in the model green alga *Chlamydomonas reinhardtii* that each EF is best suited to a specific domain of energetic fluctuation periodicity for which it can support the cell’s energetic needs, which we term bandwidth. By systematically exposing cells to a range of light periodicities, we show that while cyclic EF has a large bandwidth, pseudo-cyclic EF (PCEF) can only sustain the cell’s energetic needs for fast light fluctuations, and that the interplay between the chloroplast and the mitochondria (CMEF) has a limited bandwidth. We further show that the bandwidths of PCEF and CMEF, specialized for dynamic lights, are related to their capacity to generate ATP and protect the photosynthetic apparatus. Finally, we show that in wild-type cells, the activity level of PCEF matches its bandwidth, and we propose that cells tune the relative activity of AEFs depending on the light fluctuation frequency. Our work opens an avenue of research to characterize the molecular mechanisms that can sustain phototrophic growth in complex and dynamic energetic landscapes. It further provides a generalizable framework for understanding the physiological importance of molecular mechanisms in a dynamic environment.

## Introduction

Life depends on extracellular energy to support growth, reproduction, and survival. However, the availability of energetic vectors (i.e., organic carbon, light, reductant) is variable. It fluctuates over a wide range of timescales, with energetic overload or starvation often lasting anywhere from milliseconds to hours, days, or months (*1*, *2*). Consequently, cells must dynamically manage their cellular energy in response to these changes. Electron flows (**EFs**), active in energetic organelles such as the mitochondria or chloroplasts in eukaryotes, play a central role in cellular health by dynamically balancing cellular energy demand and supply (*3*). EFs dynamically poise ATP and redox levels in the cell by coupling electron transport with proton translocation across membranes, hence maintaining the bioenergetic homeostasis. Impairments of EFs strongly affect bioenergetic homeostasis and have been linked with aging (*4*) and multiple human diseases (*5–7*), including chronic fatigue (*8*) or long COVID-19 (*9*), and cause reduced fitness across all taxa, including yeast (*10*), nematodes (*11*), plants (*12*, *13*), or algae (*14*, *15*). While EFs’ flexibility has been proposed to be critical in modulating the cell’s bioenergetic homeostasis in response to variable energetic availability, how this flexibility is managed remains unclear.

Photosynthetic organisms, like plants or algae, are particularly subject to dynamic energy input since multiple natural factors like canopy movement, clouds, or water mixing dynamically change the amount of light energy available on timescales ranging from milliseconds to hours (*2*, *16*). Light intensity variations span a large magnitude of intensities from low, limiting light to intense, over-saturating light (*17*). In the chloroplasts of plants and eukaryotic algae, absorbed light energy is used by photosystems (**PS**) I and II to generate a linear electron flow (**LEF**) that produces NADPH and drives ATP synthesis by producing a trans-thylakoidal proton gradient (**Fig. 1A**), both of which support the energetic needs of CO_2_ conversion to biomass. However, LEF cannot sustain the cell’s energy by itself (*12*, *15*), and alternative electron flows (**AEF**) are required to balance the ATP needs and control photosynthetic electron transport by translocating additional protons across energized membranes in response to light (**Fig. 1A**). Those include (*i*) cyclic electron flow (**CEF**), which recycles reductant around PSI (*18*), (*ii*) pseudo-cyclic electron flow (**PCEF**), which reduces O_2_ into H_2_O downstream of PSI (*19–21*) and (*iii*) chloroplast to mitochondrial electron flow (**CMEF**) which involves the translocation of stromal reductant to the mitochondria (*14*, *22–24*) (**Fig. 1A**). In the model green microalgae *Chlamydomonas reinhardtii* (*Chlamydomonas* hereafter), CEF, PCEF, and CMEF are the main AEFs sustaining photosynthesis under continuous illumination (*15*). The main pathway of CEF is regulated by the proton gradient regulation-like 1 (**PGRL1**; **Fig. 1A**) (*25*), and PCEF is mostly mediated by flavodiiron proteins (**FLVs**; **Fig. 1A**)(*20*). While the molecular mechanisms involved in redox exchange between the chloroplast and the mitochondria for CMEF remain to be discovered (*26*, *27*), the majority of CMEF is mediated through the respiratory complex III (*15*) (**Fig. 1A**). Each pathway is involved in balancing the bioenergetic homeostasis in response to a dark-to-light transition.

**Figure 1.**
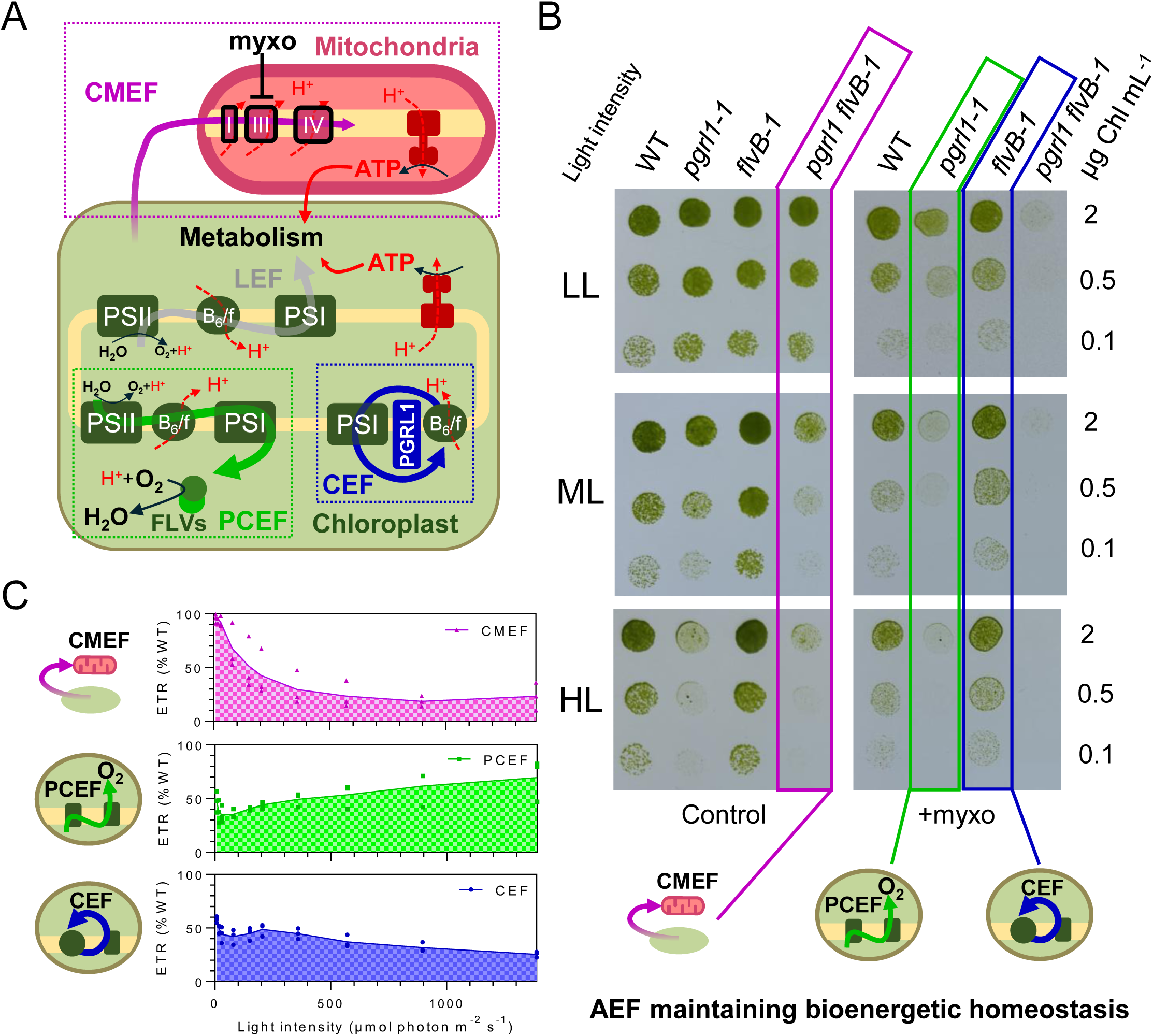
Each AEF’s capacity to maintain algal bioenergetic homeostasis in response to light widely differs. (**A)** Schematic representation of the major photosynthetic electron transport systems in the green microalgae *Chlamydomonas*. Linear electron flow (LEF, gray arrow) drives the production of NADPH and generates a trans-thylakoidal proton (H□) gradient utilized for ATP synthesis. Both NADPH and ATP fuel cell’s metabolism. Cyclic electron flow (CEF, blue arrow) and pseudo-cyclic electron flow (PCEF, green arrow) rebalance ATP generation and NADPH amounts within the chloroplast. The chloroplast-to-mitochondria electron flow (CMEF, purple arrow) allows the transformation of chloroplastic reductant into ATP mostly via the complex III/IV pathway. (**B**) Growth tests for *pgrl1, flvB, pgrl1 flvB* double mutant, and their corresponding control strains (WT) in the presence (+myxo) or absence (control) of the mitochondrial complex III inhibitor myxothiazol (2.5 *µ*M final concentration). Cells were spotted at various initial chlorophyll concentration on plates containing minimal medium at pH 7.2 and grown under continuous low light (LL, 25 *µ*mol photon m^−2^ s^−1^ for 6 days), medium light (ML, 50 *µ*mol photon m^−2^ s^−1^ for 6 days), or high light (HL, 100 *µ*mol photon m^−2^ s^−1^ for 4 days) in air enriched at 2% CO_2_. Spots shown are representative of n=10 biologically independent experiments. (**C**) Relative PSII electron transport rate (ETR) sustained by each individual AEF after 1 minute of illumination at different light intensities. Plain lines show the average, and dots show single replicates (*n =* 3 biologically independent samples).

In low-light-acclimated cells, CEF and PCEF activity peak in the first few seconds before reaching steady-state levels (*20*, *28*, *29*); while CMEF dynamic activity has not been assessed, recent measurements suggest it generates about 30% of the ATP required for CO_2_ fixation 10 minutes following a dark-to-light transition (*15*). Mutants impaired in CEF have been reported to be sensitive to high light (**HL**) and some fluctuating light (*30*, *31*). PCEF mutants are particularly affected during dark-to-light transition (*20*, *31*, *32*), and CMEF has been proposed to mostly act as a long-term electron valve in response to HL (*33*). While photosynthetic regulations under complex light changes are massively impacted by the parameters of the light shifts, like the frequency of light changes (*34–36*) or light intensities (*31*, *37*), our understanding of how the relative importance of each AEF evolves over time in dynamic environments remains limited.

This is primarily due to two factors. First, each AEF can compensate for each other (*24*, *38*), which has limited the isolation of the role of each individual AEF to maintain cellular bioenergetic homeostasis (*15*). Second, the light fluctuation regimes used in photosynthetic organisms to study AEF have largely been alternance between long phases of low light intensity and much shorter HL conditions (*20*, *31*, *39–42*). Hence, while most of our understanding has focused on how AEFs compensate (or not) for each other when low-light-acclimated cells respond to dark-to-high-light transitions, we still lack a picture of the role of each AEF across timescales in response to more complex light dynamics.

To capture the dynamic response of photosynthesis to fluctuating light, Nedbal and coworkers introduced a method based on frequency-domain analysis (43–45), in which plants are subjected to repeated periodic light fluctuations. By varying the periodicity of the light fluctuations and measuring the photosynthetic response of specific mutants, this method allows us to define domains of importance of a molecular mechanism. For example, it was used in *Arabidopsis* mutants to show that for periodicities of 10 to 60 s, the protein PsbS is crucial for the induction of the photoprotective process non-photochemical quenching (**NPQ**), while for longer periodicities, both PsbS and changes in the xanthophyll cycle contributed to the NPQ response (*36*). While powerful for characterizing the response of photosynthesis at a given time, it relies on only a few minutes of acclimation to each light fluctuation (*36*, *43*) and thus lacks the ability to integrate the importance or functional capacity of a molecular mechanism across timescales. As a result, this method is highly sensitive to the preacclimation state of the strains used.

Here, we build a framework inspired by frequency-domain analysis (*36*, *44*, *45*) to tackle the long-term role of each AEF under dynamic light conditions. We employed CRISPR-Cas9-guided mutagenesis and chemical inhibition of the mitochondrial complex III to generate mutants that rely solely on one AEF to support LEF. We first used these strains to characterise the ability of each AEF in maintaining LEF and growth under various light intensities. We then define and characterize, for each AEF, the bandwidths of light intensity fluctuation periodicities (referred hereafter simply as periodicities) for which it can sustain LEF to power at least 50% of the wild-type growth. We use this method to show that while PCEF has a bandwidth limited to periodicities shorter than a few minutes, CEF harbors a broad bandwidth across all the periodicities tested, and CMEF has a bandwidth centered around periodicities of 10 minutes. We show that the bandwidth of PCEF and CMEF reflects their capacity to sustain and protect photosynthesis in response to the first few hours of dynamic light. We further reveal that in wild-type cells acclimated to various light periodicities, the activities of PCEF and CMEF mirror their characterized bandwidth. Finally, we propose that cells regulate the mix of AEFs they use to optimize it for the periodicity of light change.

## Results

### Each AEF’s capacity to maintain energetic homeostasis depends on light intensity and exposure duration

Existing *Chlamydomonas* mutants impaired in each AEF have been previously generated in different genetic backgrounds (*20*, *23*, *25*, *33*, *46*, *47*), all of which widely differ in their genomes (*48*), transcriptomes, and photosynthetic response (*49*), making comparisons of the importance of each pathway in various environments complicated (*15*, *47*). We therefore used CRISPR Cas9-guided mutagenesis to generate multiple independent insertional mutants impaired in the accumulation of PGRL1 (***pgrl1***, impaired in CEF), FLVB (***flvB***, impaired in PCEF), or both (***pgrl1 flvB***), all in the same CC125 genetic background (**WT**) (**Fig. S1**). When two of the three main AEFs are inhibited, the third one is the only one supporting LEF in rebalancing the energetic needs of photoautotrophic cells (*15*). To isolate the capacity of each AEF in supporting growth and photosynthesis, we thus combined these mutants along with the respiratory complex III inhibitor myxothyazol (**myxo**), which inhibits the main pathway of CMEF (*50*) (**Fig. 1B** illustrates mutants/conditions for which each AEF is solely active). We first tested the capacity of each AEF to sustain growth across different light intensities, in the presence or absence of myxo (**Fig. 1B; Fig. S2**). When only CMEF was active, cells grew less than the WT under medium light (**ML**) and **HL** but grew indistinguishably from the WT under low light (**LL**) (*pgrl1 flvB* mutants, **Fig. 1B, Fig. S2**). When only CEF was active, cells grew as well as the WT for all light intensities (*flvB+myxo*, **Fig. 1B, Fig. S2-3**). However, when only PCEF was active, cells were susceptible to increasing light intensity (*pgrl1+myxo*, **Fig. 1B, Fig. S2-3**). To assess if the growth differences observed were linked with the capacity of each AEF to sustain photosynthesis, we measured the electron transport rate (**ETR**) through PSII in response to a stepwise increase in light intensity every 1 minute (**Fig. 1C; Fig. S4**). When only CMEF is active, WT levels of ETR are sustained under LL, but a marked decrease of ETR is observed with increasing light intensities (**Fig. 1C; Fig. S4**). However, when either PCEF or CEF is active, between 30% and 60% of the WT ETR is sustained for most light intensities (**Fig. 1C, Fig. S4**). Since metabolic processes like the Calvin cycle might take some time to be fully activated upon illumination, we also performed measurements of the ETR in response to stepwise light increases every 3 minutes, which showed similar results, albeit with a higher sustained ETR for each AEFs (**Fig. S5**). We conclude from these experiments that CMEF can sustain full photosynthetic activity and long-term growth only under LL. While PCEF cannot sustain the growth capacity of cells under long timescales of HL, it can sustain growth under continuous LL, suggesting that PCEF activity can be sustained at low levels for long periods of times. Furthermore, while early literature on the *Chlamydomonas pgrl1* mutant had not reported a growth defect under continuous HL (*30*, *51*), likely due to compensation by CMEF and PCEF, our results show that CEF operates best at long timescales to sustain WT levels of photosynthesis.

### PCEF and CMEF bandwidths are specialized for dynamic light conditions

Besides intensity and length of illumination, a key parameter that characterizes dynamic light intensities is how frequently light shifts from high to low intensities (*17*, *44*, *45*). To better capture the functional domain of importance for each AEF under dynamic light intensities, we used a method inspired by frequency-domain analysis, which involves measuring AEF activity and importance relative to the periodicity of a periodic light fluctuation. We first assessed the growth of strains treated and untreated with myxo in response to periodic HL/darkness shifts (**Fig. 2A**) for light periodicities ranging from one minute to four hours (**Figs. S6-S9**); continuous HL, and ML delivering the same average light energy as the fluctuating light were used as controls. As expected, when all AEF are inactive, cells do not grow for any light periodicity (*pgrl1 flvB* +myxo, **Figs. S6-S9**). We quantified the growth of mutants on solid media relative to the WT from the same plate (**Figs. S6-S9)** to estimate the relative growth loss of each mutant (**Fig. S10**). We used this data to assess the relative growth sustained when only CMEF, PCEF, or CEF are active (**Fig. 2B**). Following an analogy with frequency-domain analysis, we defined the half-growth bandwidth (hereafter bandwidth for simplicity) of an AEF as being the periodicity range for which, when it is the main AEF active, it can sustain more than 50% of the WT’s growth. We thus conclude that while CEF has a large bandwidth, sustaining growth for all periodicities tested (**Fig. 2B**), PCEF has a “high-pass” bandwidth, with substantial growth capacity sustained for periodicities lower than 10 minute (**Fig. 2B**). Surprisingly, CMEF shows a “passband” bandwidth centered around a 10 minute periodicity (**Fig. 2B**). We conclude from this experiment that while CEF can balance the cell’s energy under continuous light and for most periodicities of dynamic light, PCEF and CMEF exhibit bandwidths that make them specialized for dynamic light conditions, albeit focused on different periodicities.

**Figure 2.**
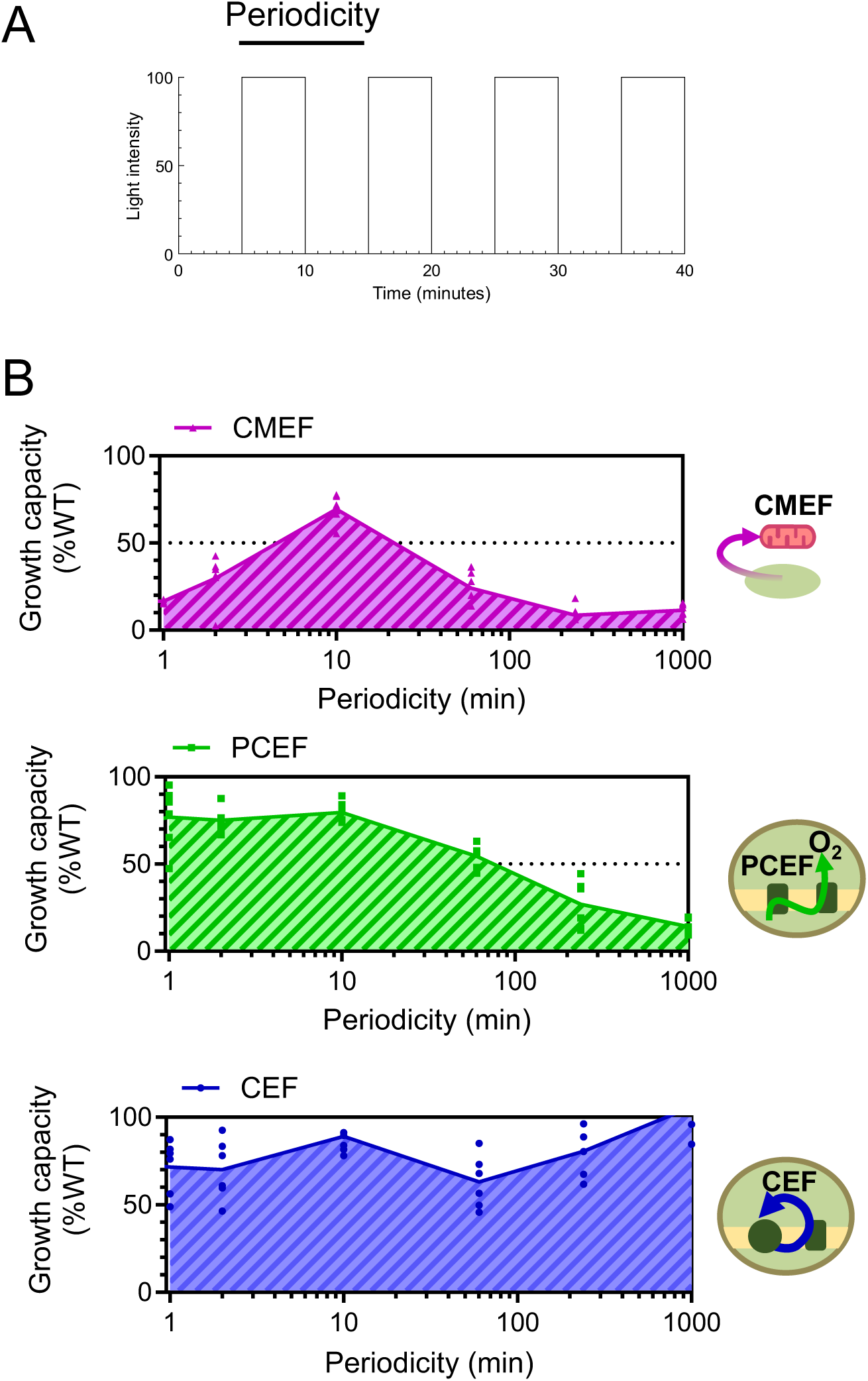
Each AEF harbors a specific bandwidth for sustaining the cell’s bioenergetic homeostasis in dynamic environments. (**A**) Illustration of the type of fluctuating light patterns used. Illustrated is a fluctuating light with a periodicity of 10 min. (**B**) Quantification of the growth capacity relative to the WT levels when only one of the main AEFs is active. Shown are greenness levels relative to the WT for the first two dilutions of spots shown in **Figs. S6-S9.** Plain lines connect averages, and dots represent the average of n = 2 technical replicates (n = 6 biologically independent samples). Note that replicates from *pgrl1-2* are not shown here as this strain showed a stronger general sensitivity to light, independent of light periodicity, which was not linked to the *pgrl1* mutation (**Figs. S6-S9**). Shown at a periodicity of 1000 minutes is continuous HL.

### PCEF and CMEF bandwidths are related to their capacity to sustain and protect photosynthesis in a dynamic light regime

AEFs contribute to the energetic balance by (i) producing ATP (ii) balancing the ATP/NADPH ratio, and (iii) establishing a proton gradient which induces mechanisms protecting the photosynthetic electron transport system (*24*, *29*). To further explore which of these mechanisms might be the most prominent for the dynamic-light-specific AEFs, we tested the ability of PCEF and CMEF to sustain LEF and maintain active PSII. We measured the ETR to assess LEF and Fv/Fm, which is a chlorophyll fluorescence parameter relative to the maximal PSII efficiency as a proxy for PSII degradation after exposure to 4 hours of HL, either continuous or fluctuating (**Fig. 3, Fig. S11**). When only CMEF was active, a slight decrease in Fv/Fm was observed for all light treatments (**Fig. 3A, Fig. S10**), suggesting that protecting PSII from inhibition does not explain the bandwidth of CMEF. Conversely, CMEF only maintained 30% of WT ETR levels under continuous HL, up to 70% for periodicities between 1 and 120 minutes (**Fig. 3A**). This suggests that the bandwidth of CMEF specialized towards dynamic conditions is dictated by the limited capacity of CMEF to produce ATP during continuous HL, and that the limited growth sustained by CMEF under 1 and 2 minutes periodicities (**Fig. 2B**) is a long-term effect. On the other hand, when only PCEF is active, the Fv/Fm is drastically reduced at HL compared to WT and increases with shorter light periodicities (**Fig. 3B, Fig. S11**). At the ETR level, only the 1 minute periodicity showed substantial ETR sustained (**Fig. 3B, Fig. S11**). This suggests that the bandwidth of PCEF is dictated by the limited capacity of PCEF to translocate protons during long periodicities of light, to both produce ATP and activate PSII protection mechanisms. When only CEF is active, both Fv/Fm and the sustained ETR showed the same trend without a clear periodicity dependence apart from a slightly higher sustained ETR at 240 minutes as compared to the other periodicities (**Fig. 3C**), suggesting that the periodicity-independent CEF role was both in sustaining LEF and protecting PSII. We conclude from this experiment that while the CMEF bandwidth mainly follows its ATP production capacity and stems from long-term imbalances, the PCEF bandwidth is dictated by both its ATP production and photoprotection capacity over shorter-term light fluctuations.

**Figure 3.**
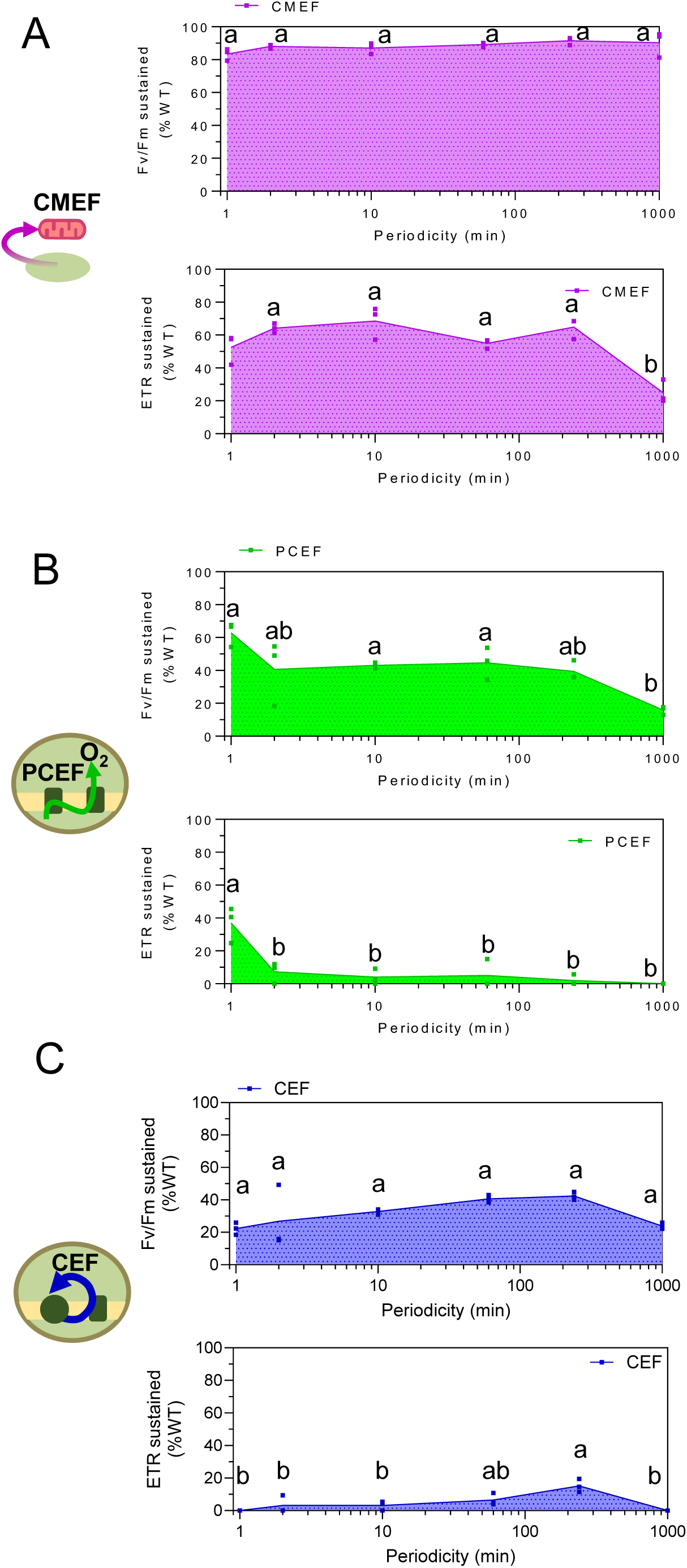
CMEF and PCEF capacity to sustain photosynthetic electron flow follows a similar behavior to their bandwidth. Maximal PSII activity (Fv/Fm; top panels) and PSII electron transport rate (ETR; bottom panels) sustained by CMEF (**A**), PCEF (**B**), or CEF (**C**) relative to the wildtype under various light periodicities after receiving photon number equivalent to 4 hours of continuous high light (100 *µ*mol photon m^−2^ s^−1^). Plain lines show the average, and dots show single replicates (*n =* 3 biologically independent samples). Shown at a periodicity of 1000 minutes is after 4 hours of continuous HL. Letters above the bars represent statistically significant differences (P value < 0.05) between conditions based on a one-way ANOVA analysis (Tukey adjusted P value); the Fv/Fm maintained thanks to CMEF did not show any statistical differences between any of the periodicities tested.

### Cells respond to the short light periodicity by activating PCEF

Given that CMEF and PCEF have bandwidths tailored for dynamic light, we hypothesized that their activity in WT cells is influenced by light periodicity to optimize AEFs’ capacities. Since both CMEF and PCEF are consuming O_2_, we used a membrane inlet mass spectrometer to measure O_2_ exchange rates in WT cells acclimated to various periodicities of light. To assess PCEF, we measure the transient O_2_ uptake rates at the onset of a dark-to-light transition (**Fig. 4, Fig. S12**), which reflects mainly PCEF (*20*). Periodicities of light shorter than 2 minutes led to an increase in the transient O_2_ uptake in response to light (**Fig. 4**); this increase was paralleled by a higher protein level of both proteins mediating PCEF for short periodicities than for long periodicities (FLVA and FLVB, **Fig. S13**). While there is no direct measurement of CMEF available, we measured respiration in the dark in an attempt to use it as a proxy for CMEF. While some trends of higher respiration were seen for periodicities of 10 minutes and 1 hour, no significant difference could be revealed between the tested periodicities (**Fig. S12**). We conclude from these experiments that WT cells acclimate to short periodicities of light by activating PCEF, which has the most relevant bandwidth.

**Figure 4.**
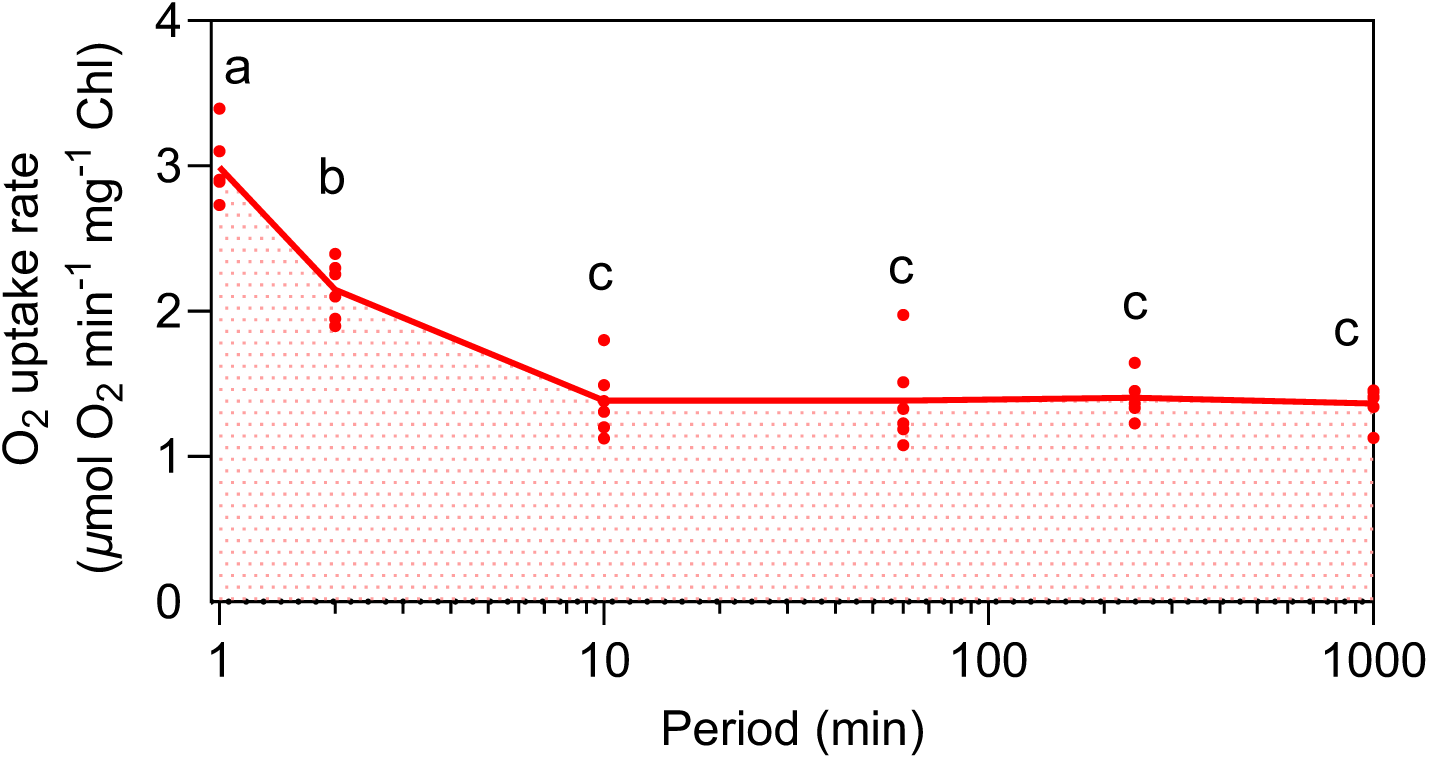
PCEF activity in WT cells parallels its bandwidth. Gas exchange rates were measured in wildtype cells after receiving a number of photons equivalent to 4 hours of continuous high light (100 *µ*mol photon m^−2^ s^−1^) for a range of light periodicities. (A, B) Gross O_2_ uptake at the onset of a dark-to-light transition (A, related to PCEF activity) or in the dark (B, related to CMEF) after cells’ acclimation to a range of light periodicities. Plain lines show the average, and dots show single replicates (*n =* 6 biologically independent samples). Shown at a periodicity of 1000 minutes is after 4 hours of continuous HL. Letters above the bars represent statistically significant differences (P value < 0.05) between conditions based on a one-way ANOVA analysis (Tukey adjusted P value).

## Discussion

Light energy availability is one of the most dynamic conditions found in the environment, with arguably the most diverse patterns, with changes happening at timescales ranging from milliseconds to days (*2*). Despite the early discovery by Garner and Allard that plant productivity depends largely on how dynamic light is (*52*), how key characteristics of light dynamics influence the molecular mechanism involved remains unclear (*17*). Here, we show that light periodicity is a major parameter defining the molecular mechanisms used in the green microalgae *Chlamydomonas* (**Figs 2-4**). Since no growth could be detected in the double *pgrl1 flvB* mutants treated with myxothyazol, we also conclude that no AEF pathway other than the ones studied here has the potential by itself to sustain growth in dynamic light conditions. Among the three main AEFs that maintain the bioenergetic homeostasis in response to light, PCEF and CMEF harbor bandwidths of physiological importance specific to short and medium periodicities of light, respectively (**Fig. 2**). We reveal that the level of activity of PCEF in WT cells acclimated to various periodicities of light parallels its bandwidth (**Fig. 4**), suggesting that photosynthetic cells sense the periodicity of light to use the most relevant AEF. Interestingly, measurements shown in **Fig. 4** are performed 15 minutes after cells are extracted from their dynamic light environment; hence, our results further suggest that the periodicity-specific acclimation involves long-term changes induced by each periodicity. For PCEF, it is clear that the decreasing amount of FLVs proteins (**Fig. S13**) might explain the low activity of PCEF under continuous light and long periodicities of light, potentially due to increased degradation of FLV proteins. However, FLVs amounts for one-minute and 10-minute periodicities are similar, suggesting a role of post-transcriptional modifications, like the thiol-based reduction of FLVA (*53*), might be involved in enhancing PCEF capacity when short periodicities are sensed. Note that a change in the protein production rate might also play a role in the lower FLV amount under HL, which we, however, did could not measure.

The bandwidths of PCEF and CMEF are qualitatively similar to their capacity to maintain LEF (**Fig. 3**) and their activation level in WT cells acclimated to each periodicity of light (**Fig. 4**). However, there are quantitative differences, which likely reflect at least two effects: (i) small imbalances created by the mutation/inhibitor add up over time and could lead to more drastic long-term effects than short-term effects as seems to be the case when only CMEF is active (**Figs. 2B; 3A**) (ii) strong imbalances created by the mutation/inhibitor might take more time to be compensated for on long timescales by metabolic adjustments, as seems to be the case when only PCEF is active (**Figs. 2B; 3B**). The quantitative differences might also stem from variation in the culture condition (liquid vs solid), which could have impacted the amount of HL perceived. Overall, similar qualitative behaviors of the capacity of either CMEF or PCEF to support various parameters relative to periodicities of light point to the bioenergetic importance of PCEF and CMEF being for short and medium periodicities of dynamic light, respectively.

Since the double *pgrl1 flvB* mutants treated with myxo did not show substantial growth or electron transport for any periodicity (**Fig. 1B; Figs. S2-11**), other molecular actors of AEF than the ones studied here likely play a minor role in the response to light fluctuations. This is in line with previous conclusions under continuous illumination (*15*) and the small fraction of electron flow that other molecular actors represent (*18*, *54*, *55*). These include the AOX pathway of CMEF (*33*), the plastid terminal oxidases (PTOX 1&2) (*55*), the plastid NAD(P)H dehydrogenase Nda2 (*56*), or Mehler reactions (*57–60*). Our work, however, does not rule out their potential roles under dynamic light conditions as fine-tuners of photosynthetic response. For example, PTOX2 has been shown to take part in rebalancing the plastoquinone redox during the dark phase of fluctuating light with a periodicity of a few minutes when the amplitude of light intensity change is wide (*61*). These secondary AEFs and their associated periodicity dependence might play a critical role in fine-tuning the activity of the main photosynthetic electron flows.

### PGRL1-controlled CEF lacks a light-periodicity or light-intensity dependence

In *Chlamydomonas*, *pgrl1* mutants are particularly sensitive to fast light fluctuations (*30*, *31*); which has been interpreted as CEF being specifically important in fluctuating light conditions. However, it is well known that in single AEF mutants, the other AEFs compensate for their loss (*24*, *30*, *38*, *62*), therefore mutants approaches are limited to defining which function of the mutated AEF cannot be compensated for. Here, by generating mutants that rely on one AEF we have isolated the bandwidth of CEF. We show that, unlike PCEF or CMEF, CEF likely does not hold a specifically high capacity of sustaining photosynthesis under fast fluctuating light conditions (**Fig. 2**), nor specifically under high light (**Fig. 1**). Our results hence suggest that CEF represents a general, versatile electron flow that can supplement LEF equally under most conditions, the other AEFs being more specialized for certain dynamic light conditions. While the PGRL1-controlled CEF might show a general broadband AEF in phototrophs, many phototrophs harbor another type of CEF, mediated by an NAD(P)H dehydrogenase-like (NDH) complex (*63*). Interestingly, in the angiosperm *Arabidopsis*, genetic ablation of the NDH-mediated CEF led to sensitivity to short periodicities of light but not to longer ones (*64*). Angiosperms have lost the FLV-mediated PCEF, which is specific to short periodicities of light; it is therefore tempting to propose that the NDH activity might have functionally evolved to complement the loss of PCEF in angiosperms.

### PCEF’s bandwidth may reflect the molecular mechanism of FLVs

Since the measurement of PCEF *in vivo* by Radmer and Kok (*65*) and the later description of the role of FLVs in PCEF (*20*, *39*), the role of PCEF has mostly been seen as a transient AEF active mainly during the first minute of dark-to-light transients. Here, we reveal that while PCEF still can sustain photosynthesis for illumination times longer than 10 minutes (**Fig. 1C**, at the end of the light curve), it cannot sustain long-term photosynthesis and growth for continuous HL (**Fig. 1B**) or long periodicities of light (**Fig. 2B**). This is likely due to degradation of FLV proteins over long timescales of HL (**Fig. S13**). High light phases, are characterized by excessive capacity to generate reducing power, and since PCEF is efficient at generating a trans-thylakoidal proton gradient (*15*, *20*, *66*) by safely producing H_2_O (*67*), and FLVs are particularly good at taking electrons (*68*), it is therefore surprising that PCEF is mainly active for short light periodicities or during light transients and cannot sustain cells’ bioenergetics during long phases of continuous saturating illumination. The catalytic mechanism of FLVs is not fully resolved (*69*), but experiments have shown that FLVs from the anaerobic bacteria *Syntrophomonas wolfei* produce H_2_O_2_ when using NADPH as an electron donor (*70*). While FLVs in phototrophs likely use ferredoxins as their main electron donor (*71*), they can also use NADPH as an electron donor (*72*). Ferredoxin is reduced quickly by PSI upon illumination, but NAPD^+^ takes time to be reduced to NADPH by the Ferredoxin-NADP reductase; hence, NADPH levels often increase with time of illumination (*73*), and steady-state levels of NADPH are positively correlated with light intensity (*74*). We thus propose that FLVs might lose their capacity over long acclimation to HL (**Fig. 1B**), or long periodicities (**Figs. 2-4**) due to H_2_O_2_ they produce in the presence of NADPH, which might lead to their degradation (**Fig. S13**). For short timescales of HL or short periodicities of light, FLVs would mostly produce H_2_O, remain active, and be a major actor in maintaining the bioenergetic balance. The H_2_O_2_ produced by FLVs might be a key factor that limits the PCEF bandwidth. If this is the case, it would also open the possibility that the cell uses it to sense differences between highly dynamic and less dynamic light environments.

### The path for electrons through CMEF is long

While the possibility of CMEF to complement chloroplast energetic deficiency was described four decades ago (*22*), its physiological relevance in wild-type plants and algae has only been recently highlighted (*13*, *15*, *47*), and the molecular mechanisms involved remain unclear. Here, we reveal that CMEF is used under dynamic light conditions, with a bandwidth centered around a periodicity of 10 minutes (**Fig. 2B**), suggesting that the CMEF pathway takes time to shuttle electrons. Amongst the main explored hypotheses on how electrons are shuttled between organelles is the oxaloacetate-malate shuttle (*26*), which directly translocates malate and oxaloacetate after their reduction or oxidation by malate dehydrogenases (MDH) in either of the organelles (*75*). Interestingly, upon a dark to light transition in *Arabidopsis*, the NAPD-dependent MDH activity oscillates for 5-10 minutes before reaching steady state (*76*). Such timing could be in accord with the bandwidth of CMEF in *Chlamydomonas*. However, it’s worth noting that *Arabidopsis,* like all angiosperms, lacks FLVs (*42*, *77*), which might impact the bandwidths of its AEF as compared to *Chlamydomonas*. Indeed, *Arabidopsis* mutants affected in the regulation of MDHs were shown to be sensitive to 1-hour light periodicity (*78*), suggesting a slightly longer bandwidth of CMEF than in *Chlamydomonas*. The relatively long periodicities of CMEF’s bandwidth might also reflect the importance of other, less direct, pathways for trafficking reductant between organelles. Such trafficking has recently been shown to involve the triose-phosphate transporter TPT3 in *Chlamydomonas*, which can transport dihydroxyacetone phosphate (DHAP) and the 3-carbon acid, 3-phosphoglycerate (3-PGA) (*79*). Since it requires DHAP to first be produced during the Calvin-Benson-Bassham cycle, it would be a good candidate for a type of CMEF that would be functional for periodicities of dozens of minutes. The systematic analysis of the bandwidth of each potential CMEF effector would reveal which CMEF route electrons take in dynamic environments.

### Acclimation to dynamic energetic environments is periodicity-dependent

Most photosynthetic organisms are subject to dynamic light intensities, including diurnal cycles to which bioenergetic mechanisms acclimate thanks to intracellular clocks (*80*). Our results suggest that photosynthetic cells can sense light shifts of sub-hour periodicities and bioenergetically acclimate to them (**Fig. 4**). Like diurnal clock mechanisms that are present across the tree of life, we propose that the periodicity-dependence of bioenergetic mechanisms, including the potential existence of sub-hour clocks, extends beyond non-photosynthetic organisms. While in some cases, like plants and algae, the periodicity comes from external input of light energy (*2*, *17*), it might also stem from dynamic internal energy demand, like in zebrafish embryos, where a 24-minute periodicity of energy dissipation is observed, which parallels the cell cycle activity (*81*). We envision that defining the bandwidth of mechanisms in response to dynamic environments extends beyond bioenergetic balance. The acclimation to dynamic changes in other biotic factors, such as temperature or nutrient availability, might require mechanisms different from those in continuous environments, each likely with its own bandwidth of physiological importance.

### Limitations of the study

By creating strains that rely mostly on one AEF, we have isolated the periodicity dependence of each pathway. However, it must be noted that in these mutants/treatment conditions, we likely overstimulate the remaining AEF, hence the characterisation done here reflects the capacity of the AEF rather than their coordinated functioning in a wild type strain. It should be noted here that since each AEF compensates for each other, mutants strains of each AEF will have altered capacity in the other AEFs (*24*, *38*), and the evaluation of coordination between each AEF in a fluctuating light context would benefit from developing a *Chlamydomonas* model building on the ones developed for *Arabidopsis* response to fluctuating lights (*43*).

We have pruposedely avoided ascribing the differences between the mutants/treatments and the wildtype to a specific molecular mechanism affected by AEF changes (i.e. NPQ, ATP generation, photosynthetic control) as we aimed to compare AEFs function focusing on their functional overlap in creating a proton gradient. Whether a specific type of photoprotection or a lack of ATP production is primarily responsible for the differences observed remains to be explored further.

While our data shows that the FLV amount is not related to PCEF activity (**Fig. 4; Fig. S13**), it is worth noting that the protein levels of FLVB in the *pgrl1* mutant are slightly lower than the WT (**Fig. S1**), and we might hence have underestimated the capacity of PCEF. Similarly, a slightly higher PGLR1 protein level is found in *flvB* mutants as compared to the WT (**Fig. S1**), we might have thus underestimated the capacity of CEF. However, we do not expect these differences to strongly affect the bandwidth of each AEF.

Finally, it is worth noting that while PGRL1 is a major regulator of CEF, it does not affect the maximal rate of CEF during the first 500ms of illumination in anoxic *Chlamydomonas* (*28*) and in *Arabidopsis*, a mutation of the PGRL1-paralog protein, PGR2, allows CEF to function in the absence of PGRL1 (*82*). Our estimations of the capacities of PCEF or CMEF might therefore be overlooking the presence of a PGRL1-independent CEF, which would be misregulated in the triple mutant. While the identity of the mediators of CEF in the green lineage is still debated (*18*, *38*, *83*), identifying the main CEF mediator will be important in further evaluating its role in sustaining the cell’s energy.

## Acknowledgments

The writing of this manuscript was assisted by Grammarly to improve readability, language, and clarity. The authors keep all accountability for the accuracy and integrity of any part of the present work. The authors thank constructive comments from all members of the Burlacot laboratory. The authors thank Çağla Aybar and Maia Lopin for helping with the initial acquisition of chlorophyll fluorescence data. We acknowledge the Chlamydomonas Resource Center, funded by the US National Science Foundation, for storing the strains described in this manuscript.

## Materials and Methods

### Strains and culture conditions

*Chlamydomonas* CC125 and *pgrl1-1* and *pgrl1-2* single mutants were already described in (*84*); all other *pgrl1*, *flvB*, and *pgrl1 flvB* double mutants were produced using CRISPR-Cas9 for this study. Each mutant is labelled as *gene(s)name-x*, where *x* is a unique identifier to differentiate genetically independent strains (obtained from a different transformation event), all information for each strain can be found in **Table S1** and **Fig. S1**. All strains developed in this study are available at the Chlamydomonas resource center (https://www.chlamycollection.org/) (their reference number can be found in **Table S1**). All strains were grown photoautotrophically in 125-mL flasks with cotton stoppers at 25°C in a buffered minimal medium (20 mM MOPS pH 7.2) under constant illumination (50 μmol photons m^−2^s^−1^, light spectrum shown in **Fig. S14A**) and shaking (120 rpm), either under ambient CO_2_ or 2% CO_2_ in air. Cells were diluted at least twice before experimentation to ensure active growing and were subsequently sampled at a concentration between 5 to 7 µg Chl mL^-1^. Throughout the manuscript, a “biologically independent experiment” refers to a replication done with a fresh new liquid culture of the strains described, seeded from a source culture, and diluted at least twice before the experiment.

For characterizations of PCEF and CMEF performed in **Fig. 4** and **Figs. S12, S13**, cells were cultured in buffered minimal medium and diluted to 1 *µ*g Chl mL^-1^ at 5 pm the day before the experiment. On the day of the experiment, 30 mL was added to a 75 cm² U-shape cell culture flask with a canted neck and placed in the treatment condition (light spectrum shown in **Fig. S14B**). Cells were treated for a total dosage of 4 hours of 100 μmol photons m^−2^s^−1^ of light under the different light treatments as follows: 4 hours of constant 100 μmol photons m^−2^s^−1^ (HL) and 8 hours of constant 50 μmol photons m^−2^s^−1^; for HL/darkness fluctuations, 6 hours of the 2h periodicity, 7.5 h of 30 minutes periodicity and 8 hours of 5 minutes, 1 minute and 30 seconds periodicities. Samples for immunodetection and gas-exchange measurements were collected at time zero and at the end of each treatment.

### Generation of mutants in the CC125 background

Multiple independent *Chlamydomonas* mutants showing targeted insertion in the *FLVB*, the *PGRL1,* or both loci were generated in the CC125 background (mt+, nit-) or single mutants background (mt+, nit-) using CRISPR/Cas-9 targeted mutagenesis (**Table S1**). Single guide RNAs (sgRNAs) (**Table S2**) were designed by CHOPCHOP (1) using version 5.6 of the *Chlamydomonas reinhardtii* genome. Ribonucleic proteins (RNPs) were prepared by duplexing the sgRNA (20% volume per volume - v/v-) and Cas-9 (IDT, Ref# 427093062) (20% v/v) with duplex buffer (IDT, Ref# 325470395) (60% v/v). Before electroporation, cells were grown in Tris-acetate-phosphate (TAP) at 22 °C under continuous illumination of 50 μmol photons m^−2^s^−1^. After treatment with autolysin for 3 hours, cells were electroporated in the presence of a hygromycin resistance cassette (2 µg mL^-1^) for the generation of single mutants, or with a paromomycin resistance (2 µg mL^-1^) cassette for the generation of *pgrl1 flvB* double mutants, and the RNP mixture. The cells were resuspended in 10 mL TAP with 40 mM sucrose and left shaking overnight in dim light (10-20 µmol photons m^-2^ s^-1^). Hygromycin-resistant or paromomycin-resistant transformants were selected on solid TAP media (2% w/v Agar) containing hygromycin (20 µg mL^-1^) or paromomycin (20 µg mL^-1^), respectively. sgRNA-targeted regions were amplified via PCR to check for full or partial insertion of the resistance cassette at the target site (**Table S3 and Fig. S1**).

### Growth test

The different *Chlamydomonas* strains were cultivated at an air level of CO_2_ under moderate light (50 *µ*mol photons m^-2^ s^-1^) at 25 °C in minimal medium at pH 7.2. Cells were collected during exponential growth and diluted in fresh minimal medium to 2, 0.5, and 0.1 µg Chl mL^-1^. Seven-microliter drops were spotted on 2.0% Agar plates of minimal medium (buffered with 20 mM MOPS pH 7.2) with or without myxothyazol (2.5 *µ*M final concentration) and exposed to air enriched with 2.5% CO_2_. Homogeneous light was supplied by LED panels (light spectrum shown in **Fig. S14C**); the different light fluctuation periodicities were obtained with power outlet controllers. The temperature was maintained at 22 °C at the level of the plates. Cells were left to grow for a duration of 5 to 20 days. At the end of the growth phase, a picture of each plate was taken. Since the chlorophyll concentration per cell did not differ between cells, cell quantity on each spot was estimated by integrating the intensity of the green channel taken over each spot and substracting the intensity of the background agar using Fiji (*85*). The final value for each spot is referred to as its “greenness” throughout the manuscript. This metric was used as a proxy for growth because it allows multiple replicates across many strains and growth conditions simultaneously. The growth capacity of each strain shown in **Fig. 2 and Fig. S3** is the relative value of greenness between the strain’s spot and the WT from the same plate and the same initial cell dilution.

### Gas exchange measurements

Gas exchange rates were measured using a membrane inlet mass spectrometer (MIMS)(*86*). Cells acclimated to the different light conditions were collected by centrifugation at 450 *× g* for 3 min and resuspended in 1.5 mL of fresh buffered minimal medium (pH 7.2) at around 25 μg chlorophyll mL^−1^. HCO_3_^−^ was subsequently added to the cell suspension (10 mM final concentration). The cell suspension was then placed in the MIMS reaction vessel (mounted with a 1 mil Teflon membrane), 100 *µ*L of ^18^O-enriched O_2_ (97% ^18^O Sigma-Aldrich, ref 490474-1L) was bubbled in the suspension. After closing the vessel, gas exchange was recorded during a dark-to-light-to-dark transition (1300 *µ*mol photons m^−2^ s^−1^; green LEDs picked at 525 nm; Luminus reference PT-121-G-L11-MPK). Light intensity was set to be saturating using weakly absorbed green light to improve homogeneity. As previously described, gross and net O_2_ exchange were calculated using the MIMS analysis software (*86*). Final chlorophyll concentration was measured at the end of the experiment.

### Chlorophyll fluorescence

To estimate the electron transport rate through PSII in **Fig. 1**, chlorophyll fluorescence was measured using a PAM fluorometer (Dual-PAM 100, Walz GmbH, Germany) with the red measuring head. Detection pulses (10 µmol photons m^-2^ s^-1^ blue light) were supplied to measure fluorescence (F) at a frequency of 20 Hz. Red saturating flashes (8000 µmol photons m^-2^ s^-1^, 300 ms, 620 nm) were delivered to measure *F*M (maximal fluorescence yield in the dark-acclimated samples) and *F*M′ (upon actinic light exposure). Fluorescence emission was detected using a long-pass filter (>700 nm). The operating yield of photosystem II (Y(II)) was calculated as (*F*M′-*F*)/*F*M′, and the relative electron transport rate (ETR) was calculated as Y(II) x I x *a* where I is the actinic light intensity and *a* is a factor correcting for the incident light non absorbed by PSII (since we could not measure *a*, here we used the machine’s default value a = 0.84, ETR calculated thus represent relative values). The Chl concentration of the sample was ∼5–8 µg Chl ml^-1,^ and the 1 mL sample was bubbled with a custom gas mixture (5% CO_2_, 20% O_2_, 75% N_2_) at a flux of 15 cm^3^ min^-1^ for proper control of the gas concentrations of the sample throughout the entire experiment duration. Cells were dark-adapted for ten minutes before starting measurements for light intensity response or light fluctuation treatment, respectively. Light response curves were obtained by incrementally increasing the actinic light intensity every 1 or 3 minutes (**Fig. 1, Figs. S4-5**).

To estimate the electron transport rate through PSII in **Fig. 3**, chlorophyll fluorescence was measured using a PAM camera fluorometer (Maxi-PAM, Walz GmbH, Germany) with red measuring LEDs. Cells were grown in flasks and transferred to 24-well plates for the light treatment, after which they were put under the camera for measurement. Cells were dark-adapted for five minutes prior to measurement. Detection pulses (10 µmol photons m^-2^ s^-1^ red light) were supplied to measure fluorescence (F) at 2 Hz. Red saturating flashes (8000 µmol photons m^-2^ s^-1^, 300 ms, 620 nm) were delivered to measure *F*M (maximal fluorescence yield in the dark-acclimated samples) and *F*M′ (upon actinic light exposure). Fluorescence emission was detected using a long-pass filter (>700 nm). The operating yield of photosystem II (Y(II)) was calculated as (*F*M′-*F*)/*F*M′ and was measured after three minutes of 12 µmol photons m^-2^s^-1^. The Chl concentration of the sample was between 3 to 5 µg Chl ml^-1,^ and the sample was spiked with 0.5 mM bicarbonate (final concentration) to avoid carbon limitation during the measurement.

### Protein extraction, SDS PAGE, and Immunoblotting

Cells were collected by centrifugation at 1000 g for 5 minutes at room temperature, and cell pellets were stored at -80. Total protein was extracted by homogenizing the cell pellet in a solution comprising 1% SDS, 50 mM DTT, 1 mM Aminocaproic acid, and 1 mM benzamidine hydrochloride, with buffering by 60 mM Tris (pH 6.8) and heating at 100°C for one minute. The resulting cell lysate was centrifuged (10000 g; 5 minutes), and the supernatant was incubated with acetone (80% final concentration) at -20°C for 1 hour. Proteins were subsequently pelleted by centrifugation (5000 g; 5 minutes) and resuspended in a storage buffer containing 1% SDS, with 60 mM Tris (pH 6.8). Protein concentration was estimated utilizing the Microplate BCA protein assay Kit (compatible with reducing agents, Thermo Scientific), with bovine serum albumin as the standard. For subsequent analysis, protein extracts (10 µg protein) were loaded onto a 12% SDS-PAGE gel, running for 1 hour at 150V in Tris-glycine SDS buffer alongside the Precision Plus Protein Dual Color Standards molecular ladder (Biorad). The proteins were then transferred to a nitrocellulose membrane using a semi-dry transfer system (Transblot turbo, Biorad). Immunoblot detection featured antibodies against FlvB (1:1000)(*20*), FlvA (1:500) (*20*), and PGRL1 (1:1000) (*25*), and other antibodies PsbA (1:10000) (AS05 084), PsaA (1:3000) (AS06 172), H3 Histone (1:5000) (AS10 710), AOX1 (1:10000) (AS06 152) and COXIIB (1:10000) (AS06 151) were procured from Agrisera (https://www.agrisera.com/). An anti-rabbit HRP-conjugated antibody (1:25000) (AS09 602) was employed as a secondary antibody for immunodetection.

## Accession numbers

Genes studied in this article can be found on https://phytozome-next.jgi.doe.gov/ under the loci Cre12.g531900 (*FLVA*), Cre16.g691800 (*FLVB*), and Cre07.g340200 (*PGRL1*).

## Supplemental Tables

**Table S1:** List of mutants generated in this study, their CC-number at the Chlamydomonas collection (U. Minnesota), and sgRNAs used to knock out target genes.

| Gene ID mutated | Strains | Genetic background | CC-number | sgRNA |
| --- | --- | --- | --- | --- |
| Cre07.g340200<br>(*PGRL1*) | *pgrl1-1* | CC125 | CC-6235 | gAB004 |
|  | *pgrl1-2* | CC125 | CC-6563 | gAB023 |
|  | *pgrl1-3* | CC125 | CC-6237 | gAB004 |
|  | *pgrl1-4* | CC125 | CC-6236 | gAB023 |
| Cre16.g691800<br>(*FLVB*) | *flvB-1* | CC125 | CC-6239 | gAB002 |
|  | *flvB-2* | CC125 | CC-6268 | gAB002 |
| Cre07.g340200/<br>Cre16.g691800<br>(*PGRL1* and *FLVB*) | *pgrl1 flvB-1* | *flvB-1* | CC-6241 | gAB002 & gAB023 |
|  | *pgrl1 flvB-2* | *pgrl1-2* | CC-6242 | gAB023 & gAB002 |
|  | *pgrl1 flvB-3* | *flvB-1* | CC-6243 | gAB002 & gAB004 |
|  | *pgrl1 flvB-4* | *pgrl1-2* | CC-6244 | gAB023 & gAB002 |

**Table S2:**
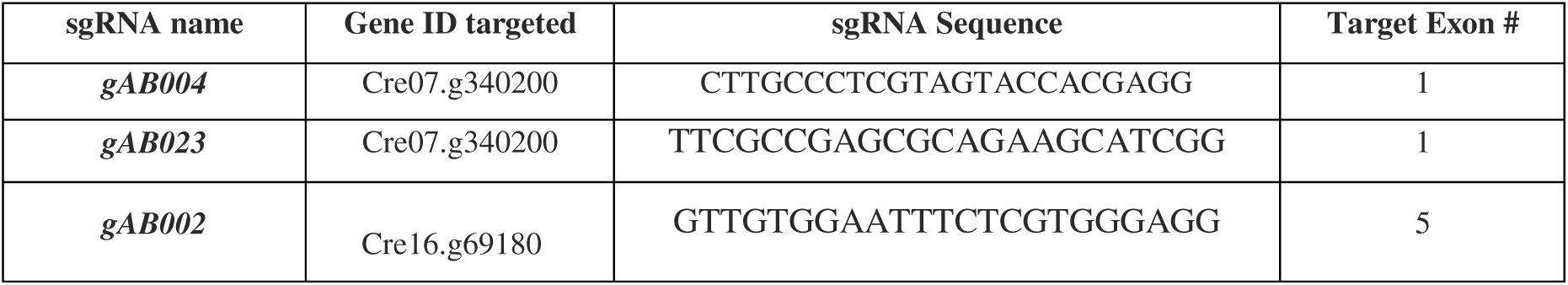
sgRNAs to knockout target genes.

**Table S3:**
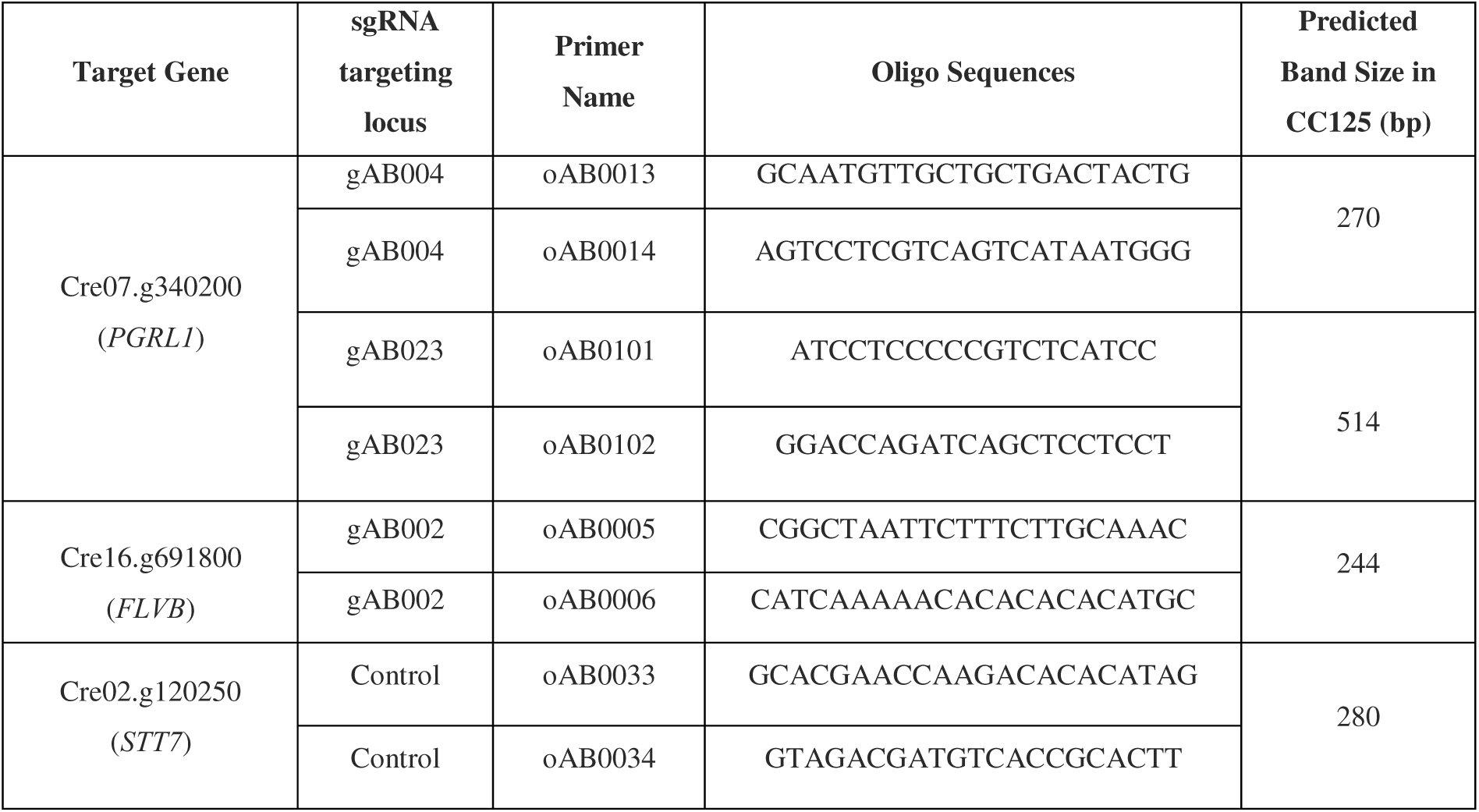
Primer pair sequences for genotyping mutants.

**Table S4:**
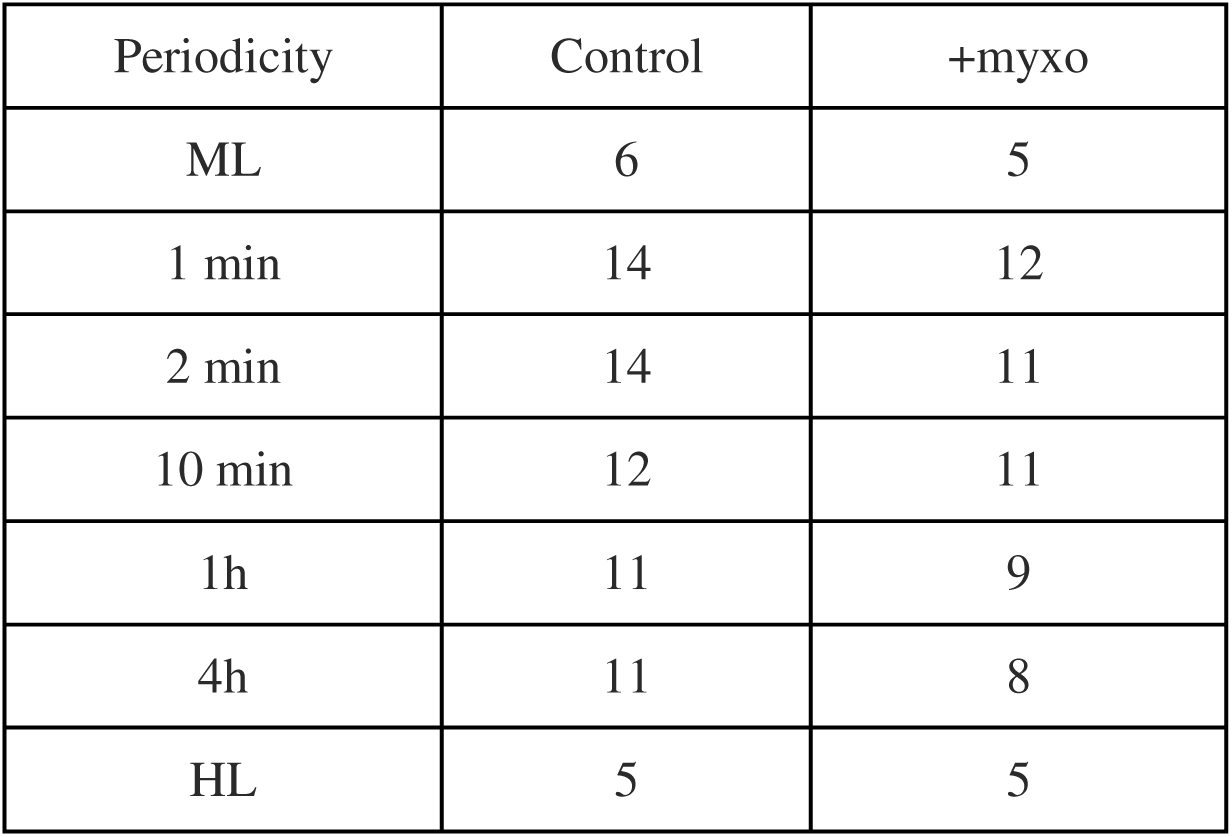
Growth time for spot tests shown in Fig. S4 and S5 (days)

**Table S5:**
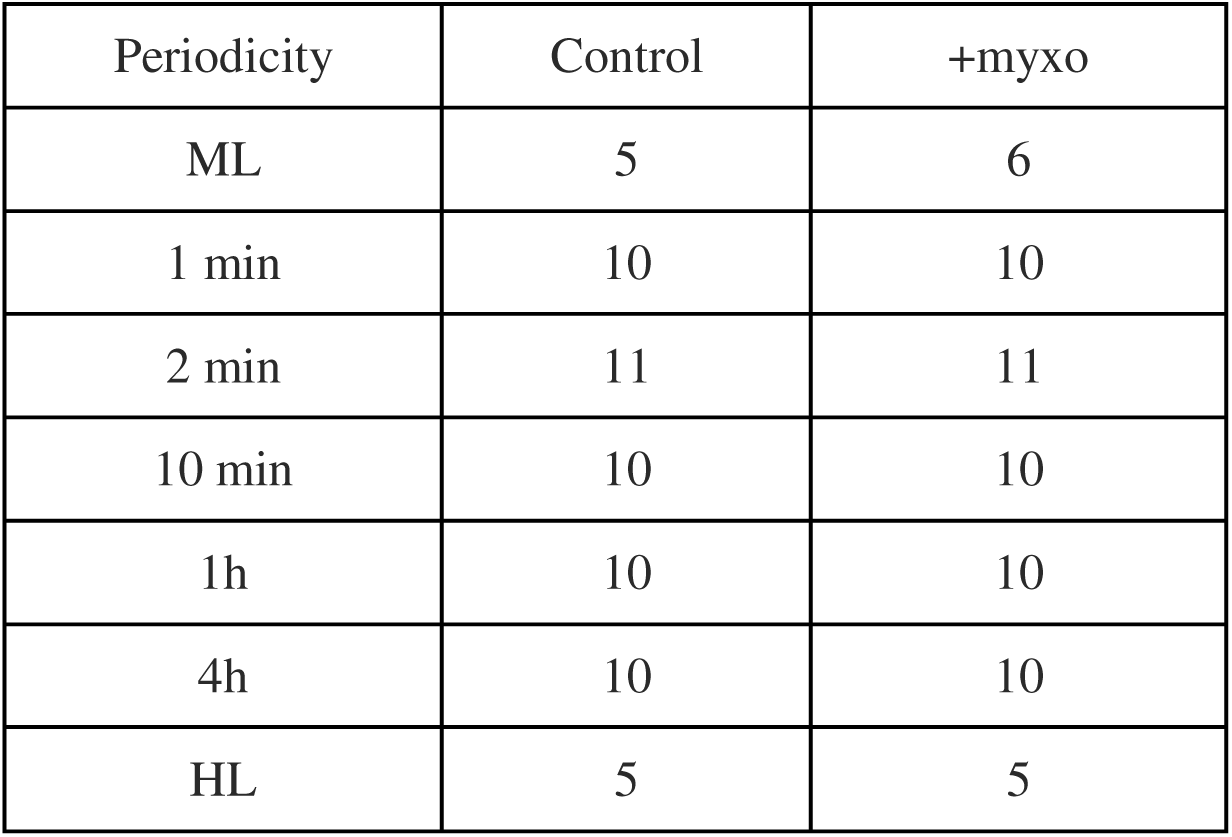
Growth time for spot tests shown in Fig. S6 and S7 (days)

| Periodicity | Control | +myxo |
| --- | --- | --- |
| ML | 5 | 6 |
| 1 min | 10 | 10 |
| 2 min | 11 | 11 |
| 10 min | 10 | 10 |
| 1h | 10 | 10 |
| 4h | 10 | 10 |
| HL | 5 | 5 |

## References

1. D. C. Wallace, Bioenergetics, the origins of complexity, and the ascent of man. Proc. Natl. Acad. Sci. 107, 8947–8953 (2010).

2. R. A. Slattery, B. J. Walker, A. P. M. Weber, D. R. Ort, The impacts of fluctuating light on crop performance. Plant Physiol. 176, 990–1003 (2017).

3. R. Putti, R. Sica, V. Migliaccio, L. Lionetti, Diet impact on mitochondrial bioenergetics and dynamics. Front. Physiol. 6 (2015).

4. A. Bratic, N.-G. Larsson, The role of mitochondria in aging. J. Clin. Invest. 123, 951–957 (2013).

5. S. B. Vafai, V. K. Mootha, Mitochondrial disorders as windows into an ancient organelle. Nature 491, 374–383 (2012).

6. D. C. Chan, Mitochondrial Dynamics and Its Involvement in Disease. Annu. Rev. Pathol. Mech. Dis. 15, 235–259 (2020).

7. E. Fernandez-Vizarra, M. Zeviani, Mitochondrial disorders of the OXPHOS system. FEBS Lett. 595, 1062–1106 (2021).

8. C. Tomas, A. Brown, V. Strassheim, J. Elson, J. Newton, P. Manning, Cellular bioenergetics is impaired in patients with chronic fatigue syndrome. PLOS ONE 12, e0186802 (2017).

9. B. D. Paul, M. D. Lemle, A. L. Komaroff, S. H. Snyder, Redox imbalance links COVID-19 and myalgic encephalomyelitis/chronic fatigue syndrome. Proc. Natl. Acad. Sci. 118, e2024358118 (2021).

10. A. Zuin, N. Gabrielli, I. A. Calvo, S. García-Santamarina, K.-L. Hoe, D. U. Kim, H.-O. Park, J. Hayles, J. Ayté, E. Hidalgo, Mitochondrial Dysfunction Increases Oxidative Stress and Decreases Chronological Life Span in Fission Yeast. PLOS ONE 3, e2842 (2008).

11. S. L. Rea, N. Ventura, T. E. Johnson, Relationship Between Mitochondrial Electron Transport Chain Dysfunction, Development, and Life Extension in Caenorhabditis elegans. PLOS Biol. 5, e259 (2007).

12. Y. Munekage, M. Hashimoto, C. Miyake, K.-I. Tomizawa, T. Endo, M. Tasaka, T. Shikanai, Cyclic electron flow around photosystem I is essential for photosynthesis. Nature 429, 579–582 (2004).

13. A. M. Vera-Vives, P. Novel, K. Zheng, S. Tan, M. Schwarzländer, A. Alboresi, T. Morosinotto, Mitochondrial respiration is essential for photosynthesis-dependent ATP supply of the plant cytosol. New Phytol. 243, 2175–2186 (2024).

14. P. Cardol, J. Alric, J. Girard-Bascou, F. Franck, F. A. Wollman, G. Finazzi, Impaired respiration discloses the physiological significance of state transitions in *Chlamydomonas*. Proc. Natl. Acad. Sci. U. S. A. 106, 15979–15984 (2009).

15. G. Peltier, C. Stoffel, J. Findinier, S. K. Madireddi, O. Dao, V. Epting, A. Morin, A. Grossman, Y. Li-Beisson, A. Burlacot, Alternative electron pathways of photosynthesis power green algal CO_2_ capture. *Plant Cell*, koae143 (2024).

17. N. A. Eckardt, Y. Allahverdiyeva, C. E. Alvarez, C. Büchel, A. Burlacot, T. Cardona, E. Chaloner, B. D. Engel, A. R. Grossman, D. Harris, N. Herrmann, M. Hodges, J. Kern, T. D. Kim, V. G. Maurino, C. W. Mullineaux, H. Mustila, L. Nikkanen, G. Schlau-Cohen, M. A. Tronconi, W. Wietrzynski, V. K. Yachandra, J. Yano, Lighting the way: Compelling open questions in photosynthesis research. Plant Cell 36, 3914–3943 (2024).

18. W. J. Nawrocki, B. Bailleul, D. Picot, P. Cardol, F. Rappaport, F. A. Wollman, P. Joliot, The mechanism of cyclic electron flow. Biochim. Biophys. Acta BBA - Bioenerg. 1860, 433–438 (2019).

19. Y. Helman, D. Tchernov, L. Reinhold, M. Shibata, T. Ogawa, R. Schwarz, I. Ohad, A. Kaplan, Genes encoding a-type flavoproteins are essential for photoreduction of O_2_ in cyanobacteria. Curr. Biol. 13, 230–235 (2003).

20. F. Chaux, A. Burlacot, M. Mekhalfi, P. Auroy, S. Blangy, P. Richaud, G. Peltier, Flavodiiron proteins promote fast and transient O_2_ photoreduction in Chlamydomonas. Plant Physiol. 174, 1825–1836 (2017).

21. Y. Allahverdiyeva, H. Mustila, M. Ermakova, L. Bersanini, P. Richaud, G. Ajlani, N. Battchikova, L. Cournac, E. M. Aro, Flavodiiron proteins Flv1 and Flv3 enable cyanobacterial growth and photosynthesis under fluctuating light. Proc. Natl. Acad. Sci. U. S. A. 110, 4111–4116 (2013).

22. C. Lemaire, F. A. Wollman, P. Bennoun, Restoration of phototrophic growth in a mutant of *Chlamydomonas reinhardtii* in which the chloroplast *atpB* gene of the ATP synthase has a deletion: an example of mitochondria-dependent photosynthesis. Proc. Natl. Acad. Sci. U. S. A. 85, 1344–1348 (1988).

23. P. Cardol, G. Gloire, M. Havaux, C. Remacle, R. Matagne, F. Franck, Photosynthesis and state transitions in mitochondrial mutants of *Chlamydomonas reinhardtii* affected in respiration. Plant Physiol. 133, 2010–2020 (2003).

24. A. Burlacot, Quantifying the roles of algal photosynthetic electron pathways: a milestone towards photosynthetic robustness. New Phytol. (2023).

25. D. Tolleter, B. Ghysels, J. Alric, D. Petroutsos, I. Tolstygina, D. Krawietz, T. Happe, P. Auroy, J. M. Adriano, A. Beyly, S. Cuine, J. Plet, I. M. Reiter, B. Genty, L. Cournac, M. Hippler, G. Peltier, Control of hydrogen photoproduction by the proton gradient generated by cyclic electron flow in *Chlamydomonas reinhardtii*. Plant Cell 23, 2619–2630 (2011).

26. O. Dao, F. Kuhnert, A. P. M. Weber, G. Peltier, Y. Li-Beisson, Physiological functions of malate shuttles in plants and algae. Trends Plant Sci. 27, 488–501 (2022).

27. G. Shimakawa, Y. Matsuda, A. Burlacot, Crosstalk between photosynthesis and respiration in microbes. J. Biosci. 49, 45 (2024).

28. W. J. Nawrocki, B. Bailleul, P. Cardol, F. Rappaport, F. A. Wollman, P. Joliot, Maximal cyclic electron flow rate is independent of PGRL1 in *Chlamydomonas*. Biochim. Biophys. Acta BBA - Bioenerg. 1860, 425–432 (2019).

29. A. Alboresi, M. Storti, T. Morosinotto, Balancing protection and efficiency in the regulation of photosynthetic electron transport across plant evolution. New Phytol. 221, 105–109 (2019).

30. K. V. Dang, J. Plet, D. Tolleter, M. Jokel, S. Cuine, P. Carrier, P. Auroy, P. Richaud, X. Johnson, J. Alric, Y. Allahverdiyeva, G. Peltier, Combined increases in mitochondrial cooperation and oxygen photoreduction compensate for deficiency in cyclic electron flow in *Chlamydomonas reinhardtii*. Plant Cell 26, 3036–50 (2014).

31. M. Jokel, X. Johnson, G. Peltier, E.-M. Aro, Y. Allahverdiyeva, Hunting the main player enabling *Chlamydomonas reinhardtii* growth under fluctuating light. Plant J. 94, 822–835 (2018).

32. A. Burlacot, A. Sawyer, S. Cuiné, P. Auroy-Tarrago, S. Blangy, T. Happe, G. Peltier, Flavodiiron-mediated O_2_ photoreduction links H_2_ production with CO_2_ fixation during the anaerobic induction of photosynthesis. Plant Physiol. 177, 1639–1649 (2018).

33. Y. Kaye, W. Huang, S. Clowez, S. Saroussi, A. Idoine, E. Sanz-Luque, A. R. Grossman, The mitochondrial alternative oxidase from *Chlamydomonas reinhardtii* enables survival in high light. J. Biol. Chem. 294, 1380–1395 (2019).

34. P. J. Graham, B. Nguyen, T. Burdyny, D. Sinton, A penalty on photosynthetic growth in fluctuating light. Sci. Rep. 7, 12513 (2017).

35. A. Bellan, F. Bucci, G. Perin, A. Alboresi, T. Morosinotto, Photosynthesis regulation in response to fluctuating light in the secondary endosymbiont alga *Nannochloropsis gaditana*. Plant Cell Physiol. 61, 41–52 (2020).

36. Y. Niu, D. Lazár, A. R. Holzwarth, D. M. Kramer, S. Matsubara, F. Fiorani, H. Poorter, S. D. Schrey, L. Nedbal, Plants cope with fluctuating light by frequency-dependent nonphotochemical quenching and cyclic electron transport. New Phytol. 239, 1869–1886 (2023).

37. M. Storti, A. Segalla, M. Mellon, A. Alboresi, T. Morosinotto, Regulation of electron transport is essential for photosystem I stability and plant growth. New Phytol. 228, 1316–1326 (2020).

38. D. Croteau, J. Alric, B. Bailleul, “Chapter 18 - The multiple routes of photosynthetic electron transfer in *Chlamydomonas reinhardtii*” in The Chlamydomonas Sourcebook *(Third Edition)*, A. R. Grossman, F.-A. Wollman, Eds. (Academic Press, London, 2023), pp. 591–613.

39. C. Gerotto, A. Alboresi, A. Meneghesso, M. Jokel, M. Suorsa, E.-M. Aro, T. Morosinotto, Flavodiiron proteins act as safety valve for electrons in Physcomitrella patens. Proc. Natl. Acad. Sci. 113, 12322–12327 (2016).

40. M. Storti, A. Alboresi, C. Gerotto, E.-M. Aro, G. Finazzi, T. Morosinotto, Role of cyclic and pseudo-cyclic electron transport in response to dynamic light changes in *Physcomitrella patens*. Plant Cell Environ. 42, 1590–1602 (2019).

41. M. Suorsa, S. Järvi, M. Grieco, M. Nurmi, M. Pietrzykowska, M. Rantala, S. Kangasjärvi, V. Paakkarinen, M. Tikkanen, S. Jansson, E.-M. Aro, PROTON GRADIENT REGULATION5 Is essential for proper acclimation of *Arabidopsis* photosystem I to naturally and artificially fluctuating light conditions. Plant Cell 24, 2934–2948 (2012).

42. H. Yamamoto, S. Takahashi, M. R. Badger, T. Shikanai, Artificial remodelling of alternative electron flow by flavodiiron proteins in Arabidopsis. Nat. Plants 2, 16012 (2016).

43. Y. Niu, D. Fuente, S. Matsubara, D. Lazár, L. Nedbal, Constitutive and Regulatory Responses of Arabidopsis thaliana to Harmonically Oscillating Light. Physiol. Plant. 177, e70421 (2025).

44. L. Nedbal, D. Lazár, Photosynthesis dynamics and regulation sensed in the frequency domain. Plant Physiol. 187, 646–661 (2021).

45. L. Nedbal, V. Březina, Complex Metabolic Oscillations in Plants Forced by Harmonic Irradiance. Biophys. J. 83, 2180–2189 (2002).

46. X. Johnson, J. Steinbeck, R. M. Dent, H. Takahashi, P. Richaud, S. I. Ozawa, L. Houille-Vernes, D. Petroutsos, F. Rappaport, A. R. Grossman, K. K. Niyogi, M. Hippler, J. Alric, Proton Gradient Regulation 5-mediated cyclic electron flow under ATP- or redox-limited conditions: A study of Δ*ATPase pgr5* and Δ*rbcL pgr5* mutants in the green alga *Chlamydomonas reinhardtii*. Plant Physiol. 165, 438–452 (2014).

47. A. Burlacot, O. Dao, P. Auroy, S. Cuiné, Y. Li-Beisson, G. Peltier, Alternative photosynthesis pathways drive the algal CO_2_ concentrating mechanism. Nature, doi: 10.1038/s41586-022-04662-9 (2022).

48. S. D. Gallaher, S. T. Fitz-Gibbon, A. G. Glaesener, M. Pellegrini, S. S. Merchant, Chlamydomonas Genome Resource for Laboratory Strains Reveals a Mosaic of Sequence Variation, Identifies True Strain Histories, and Enables Strain-Specific Studies. Plant Cell 27, 2335–2352 (2015).

49. X. Liu, O. Virtanen, S. D. Gallaher, W. J. Nawrocki, A. G. Glaesener, S. S. Merchant, R. Croce, Hidden diversity: Transcriptomic and photosynthetic variation among common ‘wild type’ Chlamydomonas strains. Plant J. 124, e70615 (2025).

50. G. von Jagow, T. A. Link, “Use of specific inhibitors on the mitochondrial bc1 complex” in Methods in Enzymology (Academic Press, 1986)vol. 126, pp. 253–271.

51. F. Chaux, X. Johnson, P. Auroy, A. Beyly-Adriano, I. Te, S. Cuiné, G. Peltier, PGRL1 and LHCSR3 compensate for each other in controlling photosynthesis and avoiding photosystem I photoinhibition during high light acclimation of Chlamydomonas cells. Mol. Plant 10, 216–218 (2017).

52. W. W. Garner, H. A. Allard, Effect of abnormally long and short alternations of light and darkness on growth and development of plants. J. Agric. Res. (1931).

53. M. E. Pérez-Pérez, A. Mauriès, A. Maes, N. J. Tourasse, M. Hamon, S. D. Lemaire, C. H. Marchand, The Deep Thioredoxome in Chlamydomonas reinhardtii: New Insights into Redox Regulation. Mol. Plant 10, 1107–1125 (2017).

54. J. Alric, J. Lavergne, F. Rappaport, Redox and ATP control of photosynthetic cyclic electron flow in *Chlamydomonas reinhardtii* (I) aerobic conditions. Biochim. Biophys. Acta BBA - Bioenerg. 1797, 44–51 (2010).

55. L. Houille-Vernes, F. Rappaport, F.-A. Wollman, J. Alric, X. Johnson, Plastid terminal oxidase 2 (PTOX2) is the major oxidase involved in chlororespiration in *Chlamydomonas*. Proc. Natl. Acad. Sci. 108, 20820–20825 (2011).

56. F. Jans, E. Mignolet, P. A. Houyoux, P. Cardol, B. Ghysels, S. Cuine, L. Cournac, G. Peltier, C. Remacle, F. Franck, A type II NAD(P) H dehydrogenase mediates light-independent plastoquinone reduction in the chloroplast of *Chlamydomonas*. Proc. Natl. Acad. Sci. U. S. A. 105, 20546–20551 (2008).

57. A. H. Mehler, Studies on reactions of illuminated chloroplasts. II. Stimulation and inhibition of the reaction with molecular oxygen. Arch. Biochem. Biophys. 34, 339–351 (1951).

58. A. H. Mehler, Studies on reactions of illuminated chloroplasts: I. Mechanism of the reduction of oxygen and other hill reagents. Arch. Biochem. Biophys. 33, 65–77 (1951).

59. A. H. Mehler, A. H. Brown, Studies on reactions of illuminated chloroplasts. III. Simultaneous photoproduction and consumption of oxygen studied with oxygen isotopes. Arch. Biochem. Biophys. 38, 365–370 (1952).

60. K. Asada, The water-water cycle in chloroplasts: Scavenging of active oxygens and dissipation of excess photons. Annu. Rev. Plant Physiol. Plant Mol. Biol. 50, 601–639 (1999).

61. W. J. Nawrocki, F. Buchert, P. Joliot, F. Rappaport, B. Bailleul, F.-A. Wollman, Chlororespiration controls growth under intermittent light. Plant Physiol. 179, 630–639 (2019).

62. J. Alric, X. Johnson, Alternative electron transport pathways in photosynthesis: a confluence of regulation. Curr. Opin. Plant Biol. 37, 78–86 (2017).

63. G. Peltier, E. M. Aro, T. Shikanai, NDH-1 and NDH-2 plastoquinone reductases in oxygenic photosynthesis. Annu. Rev. Plant Biol*. Vol* 67 **67**, 55–80 (2016).

64. G. Shimakawa, C. Miyake, Changing frequency of fluctuating light reveals the molecular mechanism for P700 oxidation in plant leaves. Plant Direct 2, e00073 (2018).

65. R. J. Radmer, B. Kok, Photoreduction of O_2_ pimes and replaces CO_2_ assimilation. Plant Physiol. 58, 336–340 (1976).

66. G. Shimakawa, K. Ishizaki, S. Tsukamoto, M. Tanaka, T. Sejima, C. Miyake, The Liverwort, Marchantia, drives alternative electron flow using a flavodiiron protein to protect PSI. Plant Physiol. 173, 1636–1647 (2017).

67. A. Alboresi, M. Storti, L. Cendron, T. Morosinotto, Role and regulation of class-C flavodiiron proteins in photosynthetic organisms. Biochem. J. 476, 2487–2498 (2019).

68. N. Rizzetto, E. Traverso, A. Sabia, F. Fiorin, C. Francese, L. Trainotti, T. Morosinotto, A. Alboresi, Flavodiiron protein activity outcompetes cyclic electron transport when expressed in angiosperm Nicotiana tabacum. bioRxiv [Preprint] (2025). 10.1101/2025.01.23.634494.

69. M. R. A. Blomberg, P. Ädelroth, Reduction of molecular oxygen in flavodiiron proteins - Catalytic mechanism and comparison to heme-copper oxidases. J. Inorg. Biochem. 255, 112534 (2024).

70. M. C. Martins, C. M. Alves, M. Teixeira, F. Folgosa, The flavodiiron protein from Syntrophomonas wolfei has five domains and acts both as an NADH:O2 or an NADH:H2O2 oxidoreductase. FEBS J. 291, 1275–1294 (2024).

71. P. Sétif, G. Shimakawa, A. Krieger-Liszkay, C. Miyake, Identification of the electron donor to flavodiiron proteins in Synechocystis sp. PCC 6803 by in vivo spectroscopy. Biochim. Biophys. Acta BBA - Bioenerg. 1861, 148256 (2020).

72. C. Beraldo, E. Traverso, M. Boschin, L. Cendron, T. Morosinotto, A. Alboresi, Physcomitrium patens flavodiiron proteins form heterotetrametric complexes. J. Biol. Chem. 300 (2024).

73. J. H. Jang, D. B. Kim, Y. Choi, R. Amir, D.-E. Cheong, H.-J. Chung, S.-H. Ahn, G.-J. Kim, D. W. Lee, O. R. Lee, E.-S. Kim, Real-time monitoring of stromal NADPH levels in Arabidopsis using a metagenome-derived NADPH-binding fluorescent protein. Plant Physiol. Biochem. 217, 109260 (2024).

74. K. Tanaka, G. Shimakawa, H. Tabata, S. Kusama, C. Miyake, S. Nakanishi, Quantification of NAD(P)H in cyanobacterial cells by a phenol extraction method. Photosynth. Res. 148, 57–66 (2021).

75. J. Selinski, R. Scheibe, Malate valves: old shuttles with new perspectives. Plant Biol, doi: 10.1111/plb.12869 (2018).

76. R. Scheibe, M. Stitt, Comparison of Nadp-Malate Dehydrogenase Activation, Qa Reduction and O-2 Evolution in Spinach Leaves. Plant Physiol. Biochem. (1988).

77. G. Peltier, D. Tolleter, E. Billon, L. Cournac, Auxiliary electron transport pathways in chloroplasts of microalgae. Photosynth. Res. 106, 19–31 (2010).

78. Y. Yokochi, K. Yoshida, F. Hahn, A. Miyagi, K. Wakabayashi, M. Kawai-Yamada, A. P. M. Weber, T. Hisabori, Redox regulation of NADP-malate dehydrogenase is vital for land plants under fluctuating light environment. Proc. Natl. Acad. Sci. 118, e2016903118 (2021).

79. W. Huang, A. Krishnan, A. Plett, M. Meagher, N. Linka, Y. Wang, B. Ren, J. Findinier, P. Redekop, N. Fakhimi, R. G. Kim, D. A. Karns, N. Boyle, M. C. Posewitz, A. R. Grossman, Chlamydomonas mutants lacking chloroplast TRIOSE PHOSPHATE TRANSPORTER3 are metabolically compromised and light-sensitive. *Plant Cell*, koad095 (2023).

80. D. Bell-Pedersen, V. M. Cassone, D. J. Earnest, S. S. Golden, P. E. Hardin, T. L. Thomas, M. J. Zoran, Circadian rhythms from multiple oscillators: lessons from diverse organisms. Nat. Rev. Genet. 6, 544–556 (2005).

81. J. Rodenfels, K. M. Neugebauer, J. Howard, Heat Oscillations Driven by the Embryonic Cell Cycle Reveal the Energetic Costs of Signaling. Dev. Cell 48, 646–658.e6 (2019).

82. T. Rühle, M. Dann, B. Reiter, D. Schünemann, B. Naranjo, J.-F. Penzler, T. Kleine, D. Leister, PGRL2 triggers degradation of PGR5 in the absence of PGRL1. Nat. Commun. 12, 3941 (2021).

83. T. Shikanai, Molecular Genetic Dissection of the Regulatory Network of Proton Motive Force in Chloroplasts. Plant Cell Physiol. 65, 537–550 (2024).

84. O. Dao, A. Burlacot, F. Buchert, M. Bertrand, P. Auroy, C. Stoffel, S. K. Madireddi, J. Irby, M. Hippler, G. Peltier, Y. Li-Beisson, Cyclic and pseudo-cyclic electron pathways play antagonistic roles during nitrogen deficiency in Chlamydomonas reinhardtii. Plant Physiol. 197, kiae617 (2025).

85. J. Schindelin, I. Arganda-Carreras, E. Frise, V. Kaynig, M. Longair, T. Pietzsch, S. Preibisch, C. Rueden, S. Saalfeld, B. Schmid, J.-Y. Tinevez, D. J. White, V. Hartenstein, K. Eliceiri, P. Tomancak, A. Cardona, Fiji: an open-source platform for biological-image analysis. Nat. Methods 9, 676–682 (2012).

86. A. Burlacot, F. Burlacot, Y. Li-Beisson, G. Peltier, Membrane Inlet Mass Spectrometry: A Powerful Tool for Algal Research. Front. Plant Sci. 11 (2020).

