## Supplemental figures for "Dynamic environments require photosynthetic electron flows with distinct bandwidths"

Madireddi *et al*,

Corresponding author: Adrien Burlacot


**This PDF file includes:**

Supplementary Text
Figs. S1 to S12

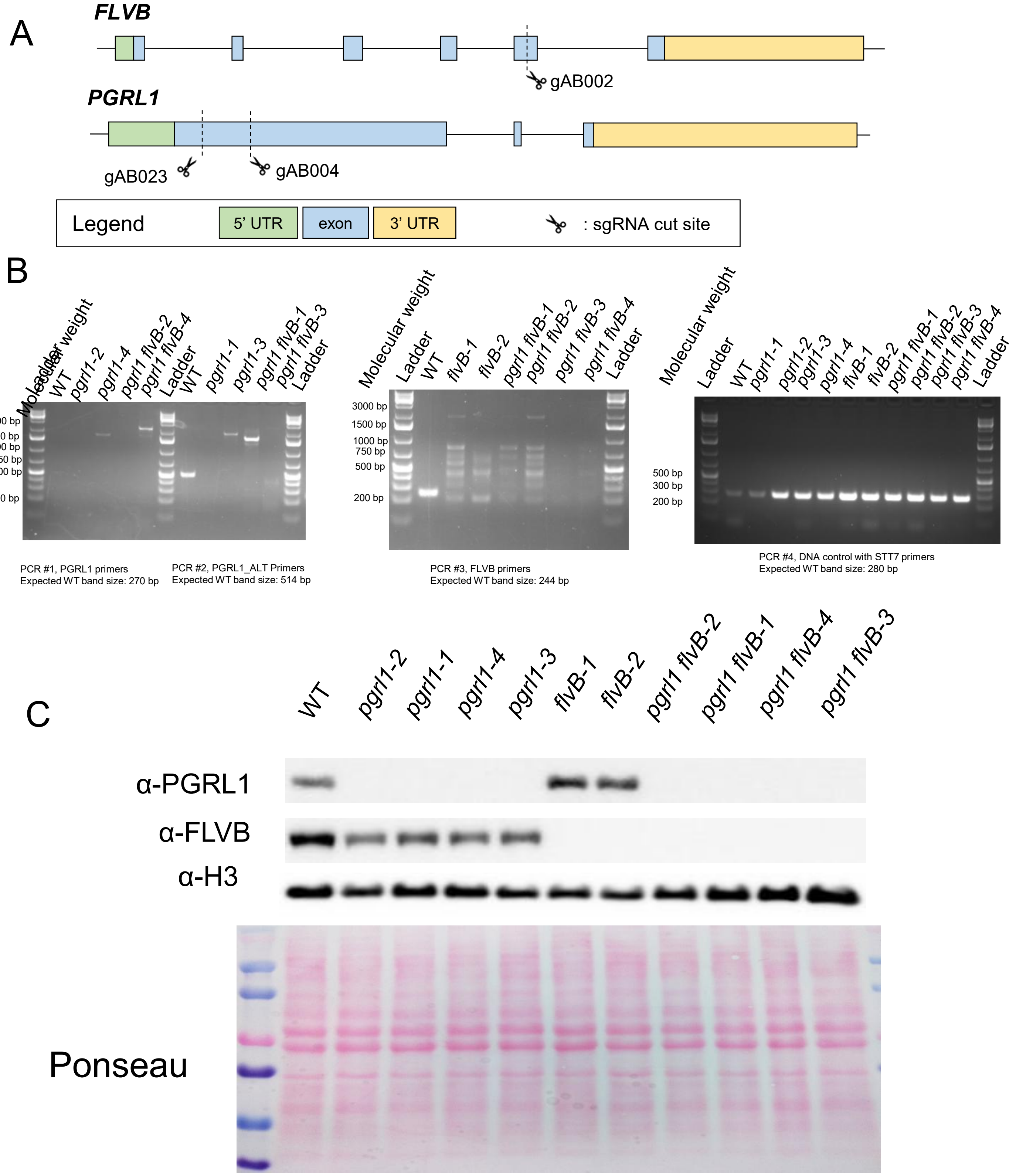


**Supplemental Figure S1. Design and characterization of new mutants carrying insertions in the *FLVB* and *PGRL1* genes. A.** Gene context of *FLVB* and *PGRL1* genes with each target cut site of sgRNA-guides used in this study. **B.** PCR verification of the insertion of the hygromycin resistance cassette in each locus of interest for all strains used in this study. **C.** Immunodetection of FLVB and PGRL1 in the strains used in this study. Immunodetection of Histone and Ponceau are shown as loading controls. Supports **Figs. 1-4**.

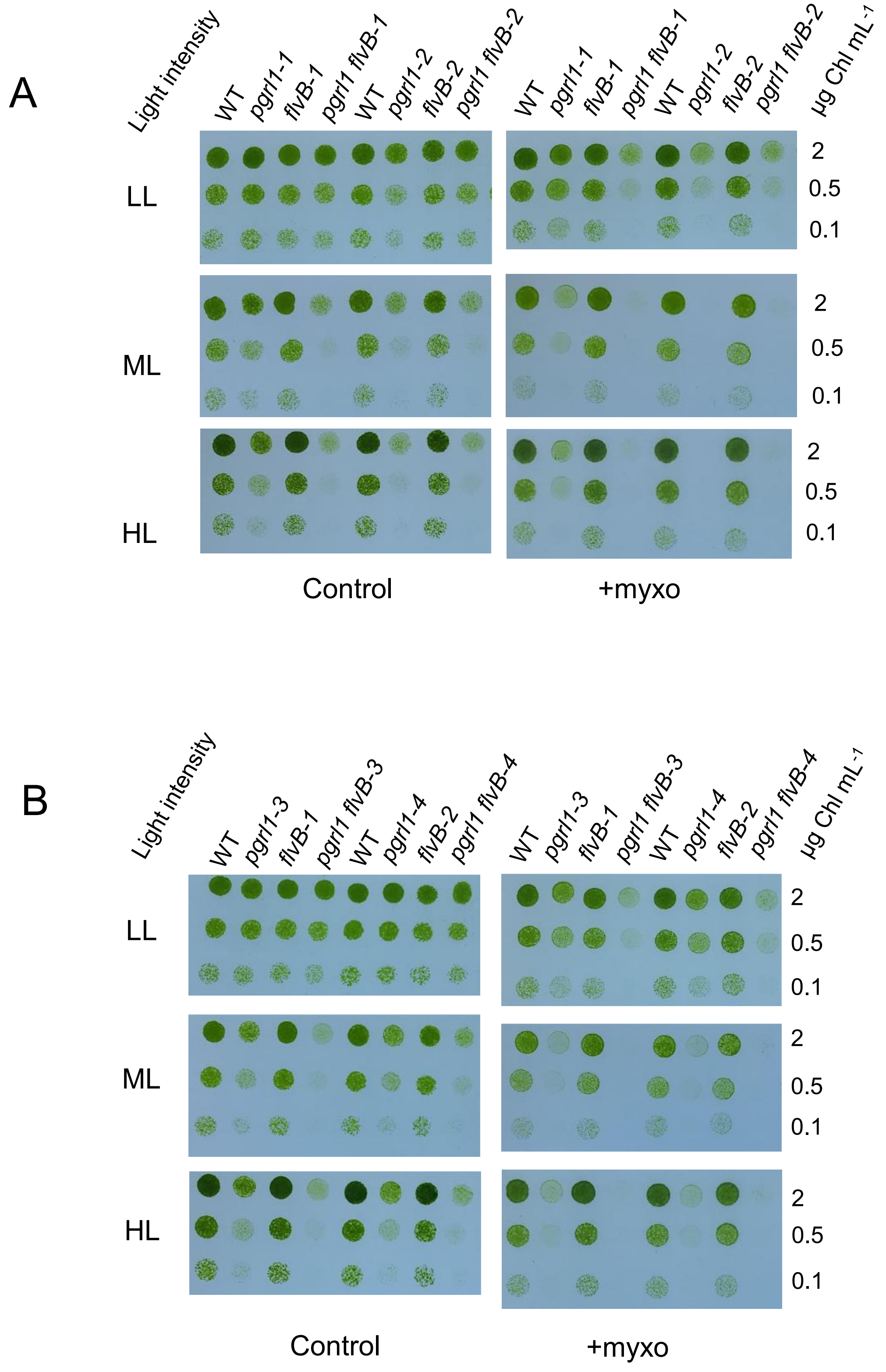


**Supplemental Figure S2. Growth test of *flvB*, *pgrl1,* and *pgrl1 flvB* mutants under various continuous light intensities.** Growth test of *flvB*, *pgrl1,* and *pgrl1 flvB* double mutants (-1 and -2 **A** and -3 and -4 **B**) and their control strain (WT) under low (25 µmol photon $m^{-2}$ $s^{-1}$, LL), medium (50 µmol photon $m^{-2}$ $s^{-1}$, ML), or high (100 µmol photon $m^{-2}$ $s^{-1}$, HL) light. Cells were spotted on plates containing minimal medium at pH 7.2 and grown under continuous illumination in the absence (left panels, Control) or presence (right panels, +myxo) of 2.5 *µ*M myxothyazol in air enriched at 2.5% $CO_2$. Cells were left to grow for 5 days (HL and ML) or 7 days (LL). Spots shown are representative of *n*=5 biologically independent experiments. Supports **Fig. 1**.

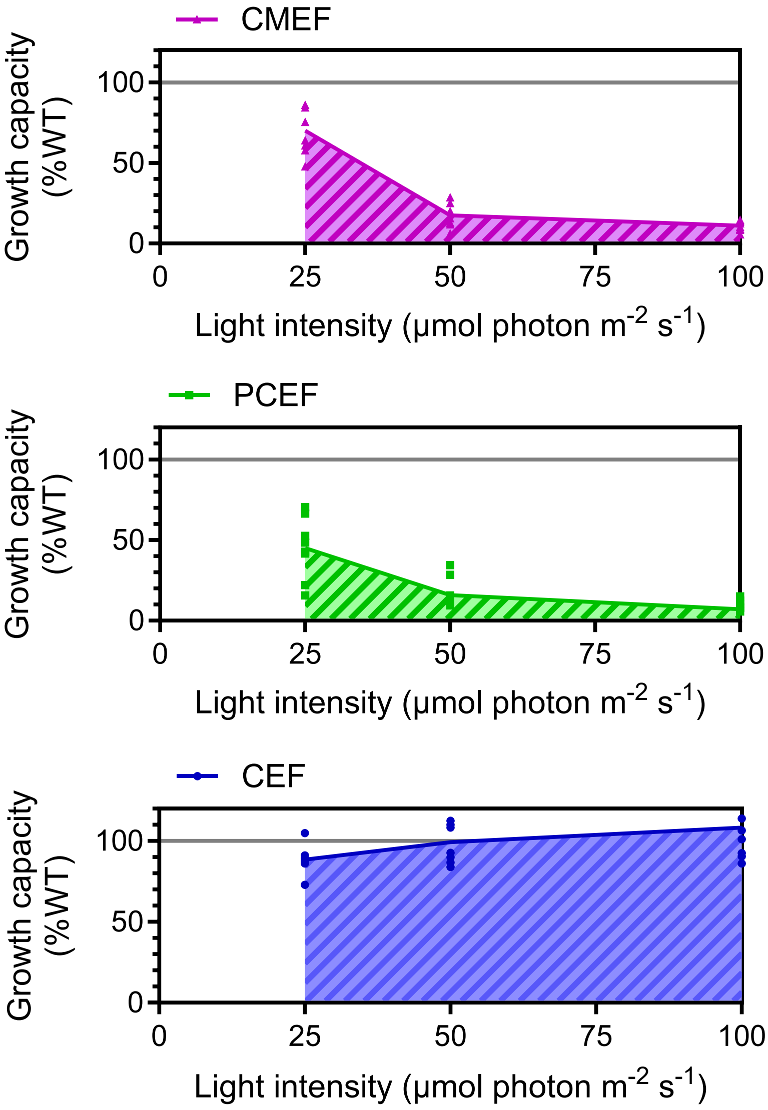


**Supplemental Figure S3. Each AEF's capacity to maintain algal growth in response to light intensity widely differs.** Quantification of the growth capacity relative to the WT levels when only one of the main AEFs is active. Shown are greenness levels relative to the WT for the first two dilutions of spots shown in **Figs. S2** for the *pgrl1 flvB* double mutants (CMEF), *pgrl1* mutants +myxo (PCEF), and *flvB* mutants +myxo **.** Plain lines connect averages, and dots represent the average of *n* = 2 technical replicates (*n* = 4 biologically independent samples)**.** Supports **Fig. 1**.

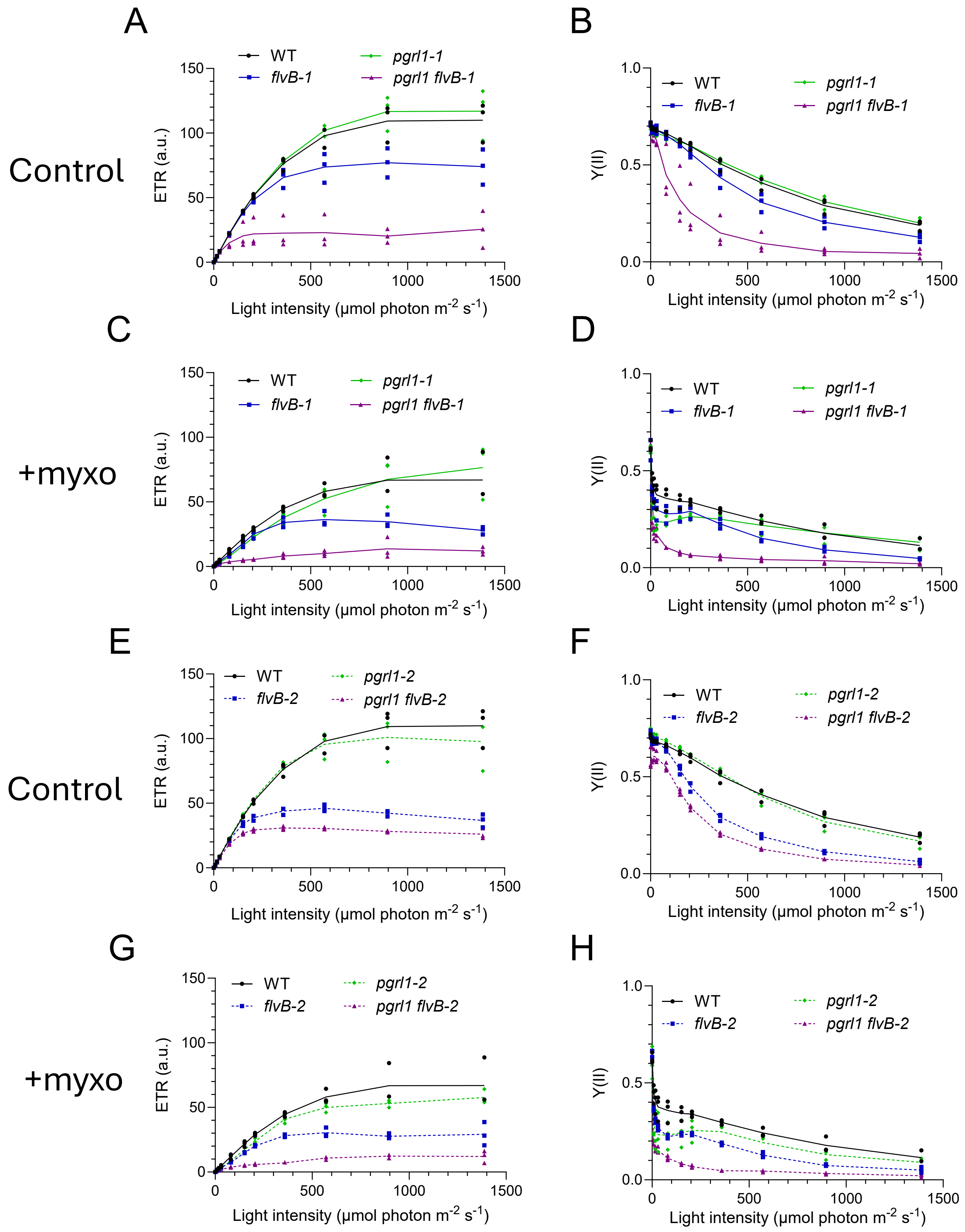


**Supplemental Figure S4. Photosynthetic activity in *pgrl1*, *flvB*, *pgrl1 flvB,* and their control strain during a stepwise increase in light intensity.** (**A, C, E, G**) PSII electron transport rate measured after 1 min of illumination at different light intensities on *pgrl1*, *flvB*, *pgrl1 flvB* mutants, and their control strain (WT) in the absence (control; **A, E**) or presence (+myxo; **C, G**) of myxothyazol (2.5 µM). (**B, D, F, H**) Corresponding PSII yield (Y(II)) for the same experiment as in **A**, **C**, **E,** and **G**, respectively. The WT shown in (**A, B**) and (**E, F**), as well as (**C, D**) and (**G, H**) are respectively the same. Lines show the average, and dots show single replicates (*n* = 3 biologically independent samples for each of the genetically independent mutants). Supports **Fig. 1**.

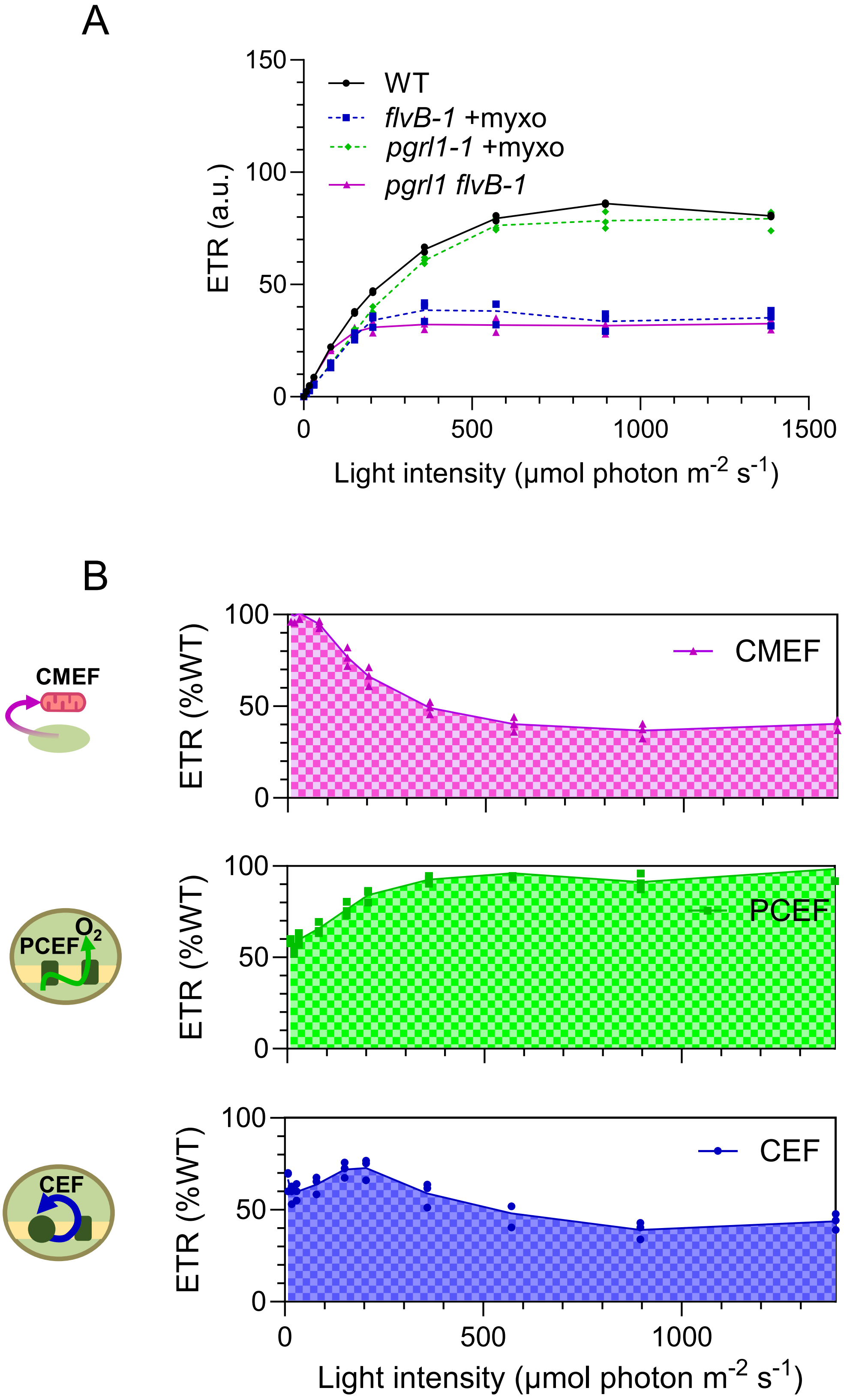


**Supplemental Figure S5. Photosynthetic activity during a stepwise increase in light intensity every 3 minutes.** (**A**) PSII electron transport rate measured after 3 min of illumination at different light intensities on *pgrl1-1*, *flvB-1*, *pgrl1 flvB-1* mutants, and their control strain (WT) in the absence (WT, *pgrl1 flvB-1*) or presence (+myxo; *pgrl1-1*, *flvB-1*) of myxothyazol (2.5 µM). (**B**) Relative PSII electron transport rate (ETR) sustained by each individual AEF after 3 minutes of illumination at different light intensities. Lines show the average, and dots show single replicates (*n* = 3 biologically independent samples for each of the genetically independent mutants). Supports **Fig. 1**.

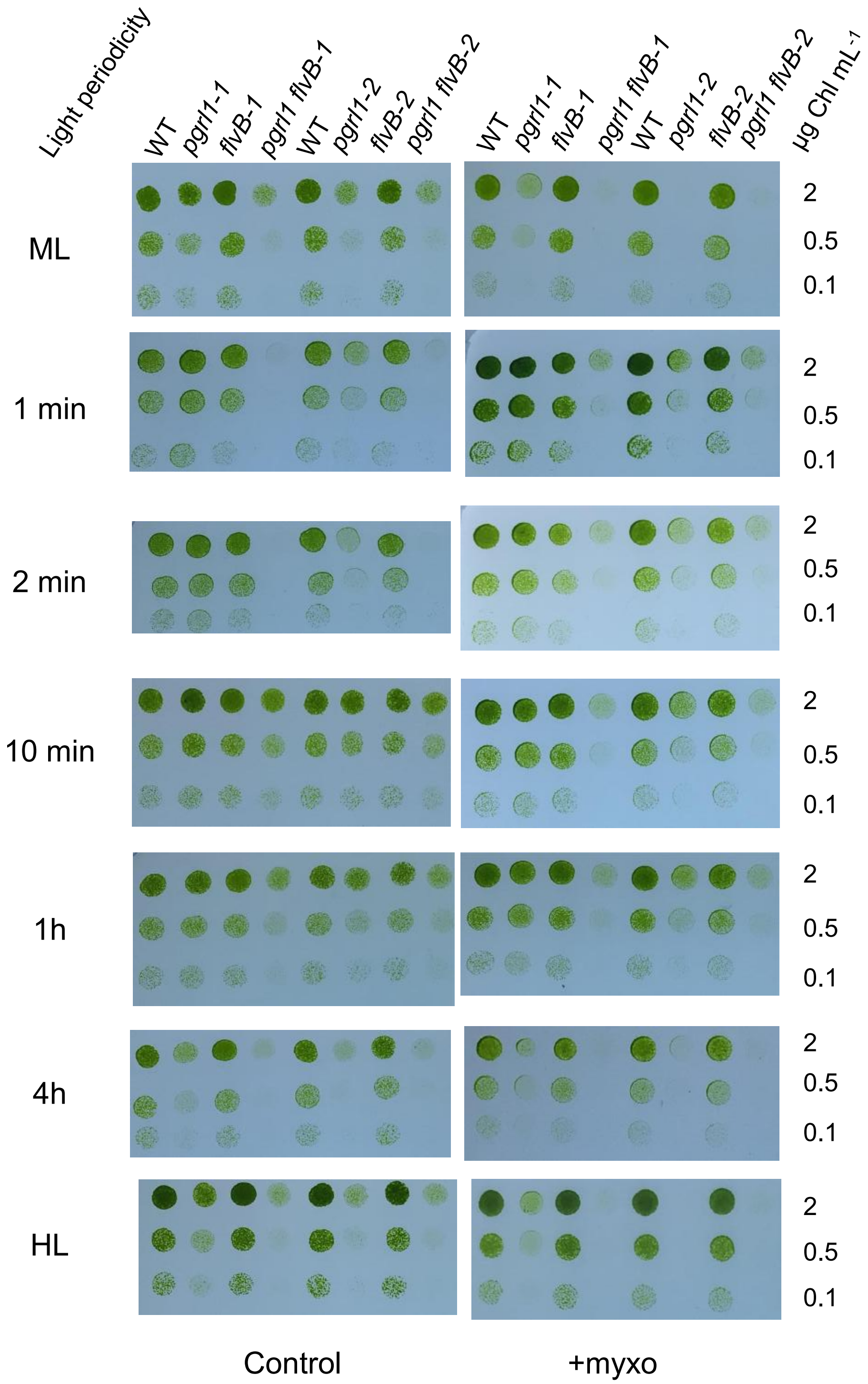


**Supplemental Figure S6. Growth test of *flvB*, *pgrl1,* and double mutants under various periodicities of light fluctuation.** Growth test of *flvB*, *pgrl1,* and *pgrl1 flvB* double mutants (-1 and -2) and their control strain under periodic darkness/high light (0/100 µmol photon $m^{-2}$ $s^{-1}$) shifts with various periodicities or continuous medium (ML, 50 µmol photon $m^{-2}$ $s^{-1}$) or high (HL, 100 µmol photon $m^{-2}$ $s^{-1}$) light. Cells were spotted on plates containing minimal medium at pH 7.2 and grown under the various illumination patterns in the absence (left panels) or presence (right panels) of 2.5 *µ*M myxothyazol in air enriched at 2.5 % $CO_2$. Cells were left to grow between 5 to 14 days (**Table S.4**). Spots shown are representative of *n*=5 biologically independent experiments. Supports **Fig. 2**.

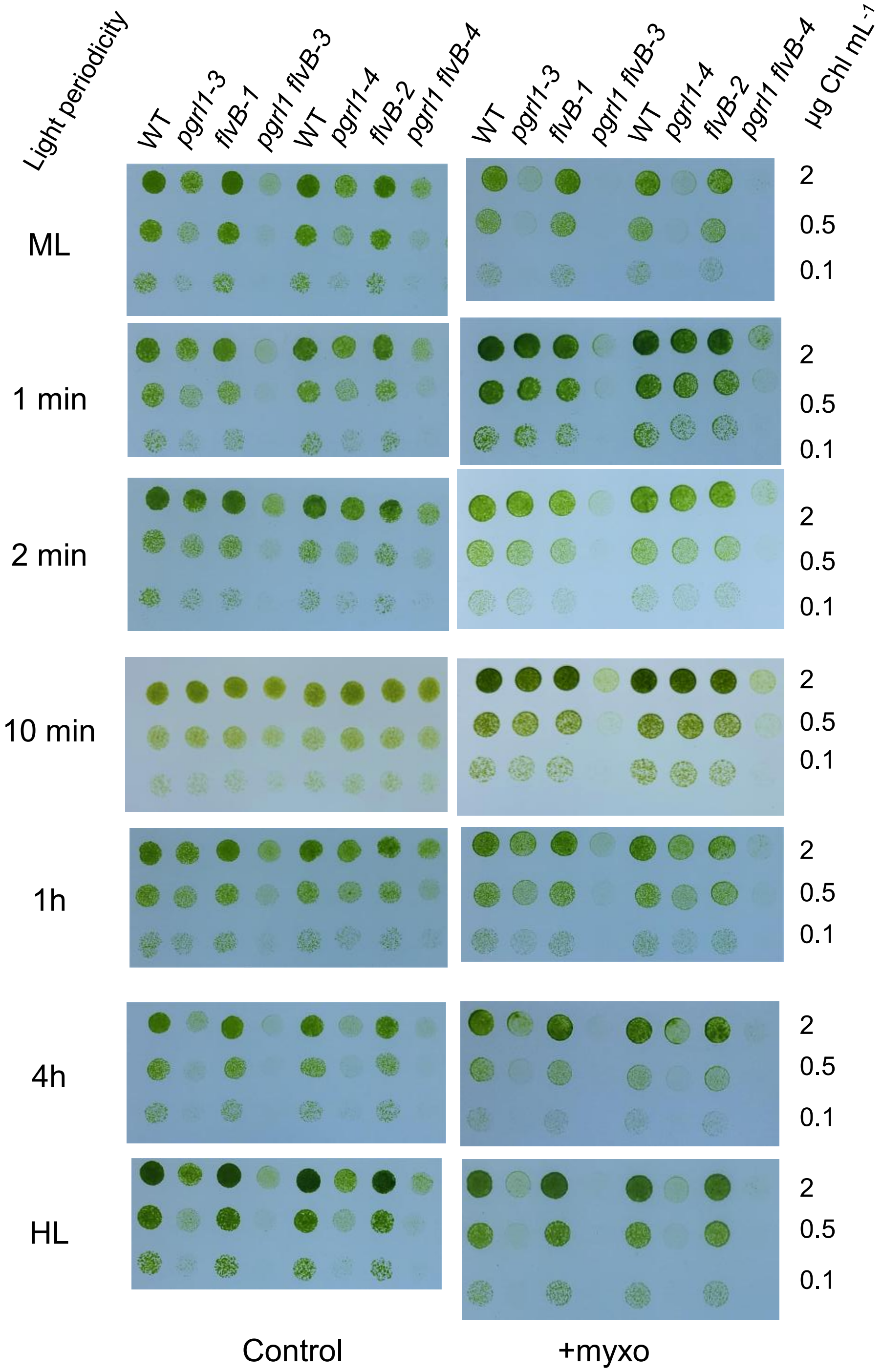


**Supplemental Figure S7. Growth test of *flvB*, *pgrl1,* and double mutants under various periodicities of light fluctuation.** Growth test of *flvB*, *pgrl1,* and *pgrl1 flvB* double mutants (-3 and -4) and their control strain under periodic darkness/high light (0/100 µmol photon $m^{-2}$ $s^{-1}$) shifts with various periodicities or continuous medium (ML, 50 µmol photon $m^{-2}$ $s^{-1}$) or high (HL, 100 µmol photon $m^{-2}$ $s^{-1}$) light. Cells were spotted on plates containing minimal medium at pH 7.2 and grown under the various illumination patterns in the absence (left panels) or presence (right panels) of 2.5 *µ*M myxothyazol in air enriched at 2.5 % $CO_2$. Cells were left to grow between 5 to 14 days (**Table S.4**). Spots shown are representative of *n*=5 biologically independent experiments. Supports **Fig. 2**.

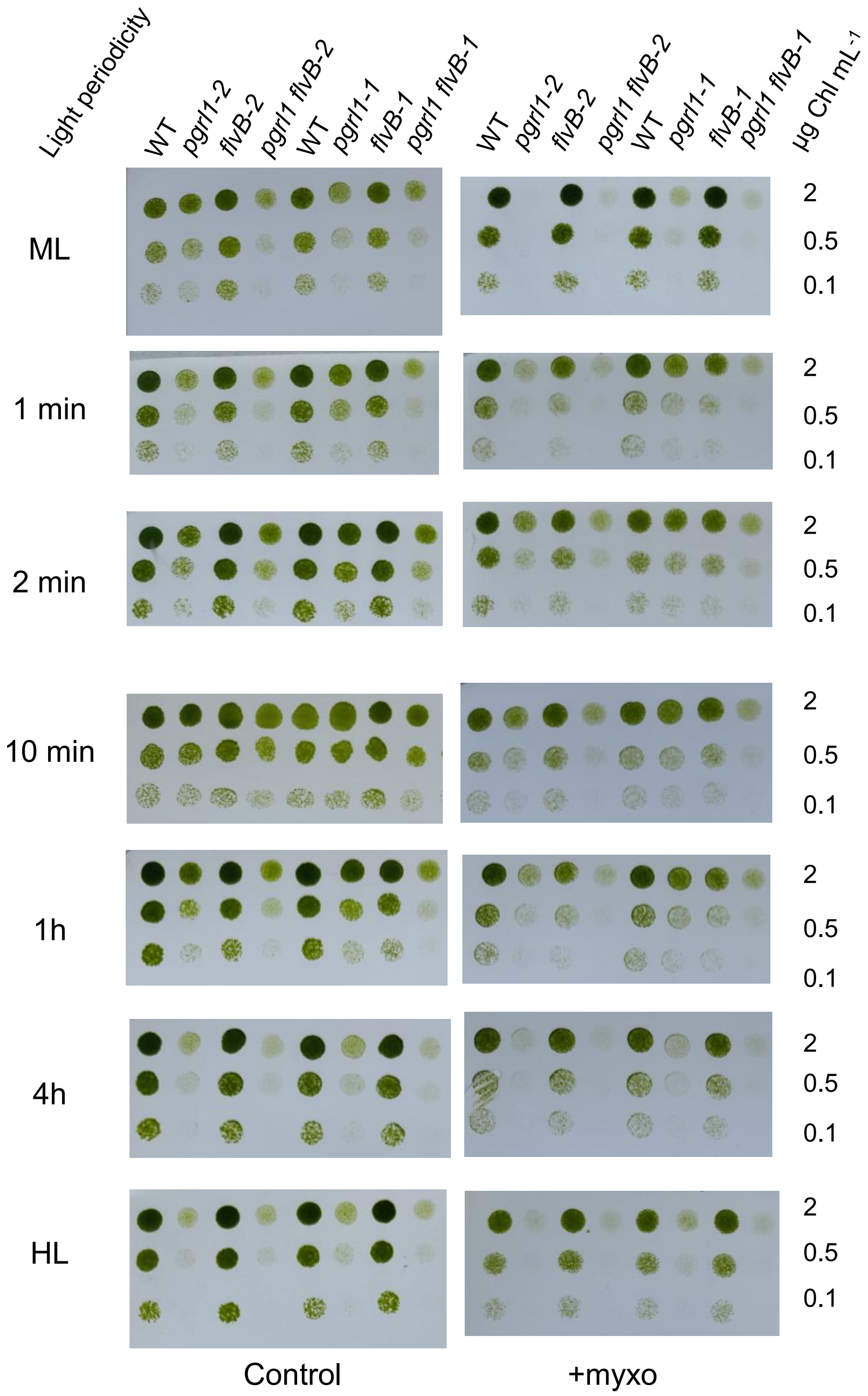


**Supplemental Figure S8. Growth test of *flvB*, *pgrl1,* and double mutants under various periodicities of light fluctuation.** Growth test of *flvB*, *pgrl1,* and *pgrl1 flvB* double mutants (-1 and -2) and their control strain under periodic darkness/high light (0/100 µmol photon $m^{-2}$ $s^{-1}$) shifts with various periodicities or continuous medium (ML, 50 µmol photon $m^{-2}$ $s^{-1}$) or high (HL, 100 µmol photon $m^{-2}$ $s^{-1}$) light. Cells were spotted on plates containing minimal medium at pH 7.2 and grown under the various illumination patterns in the absence (left panels) or presence (right panels) of 2.5 *µ*M myxothyazol in air enriched at 2.5 % $CO_2$. Cells were left to grow between 5 to 11 days (**Table S.5**). Spots shown are representative of *n*=5 biologically independent experiments. Supports **Fig. 2**.

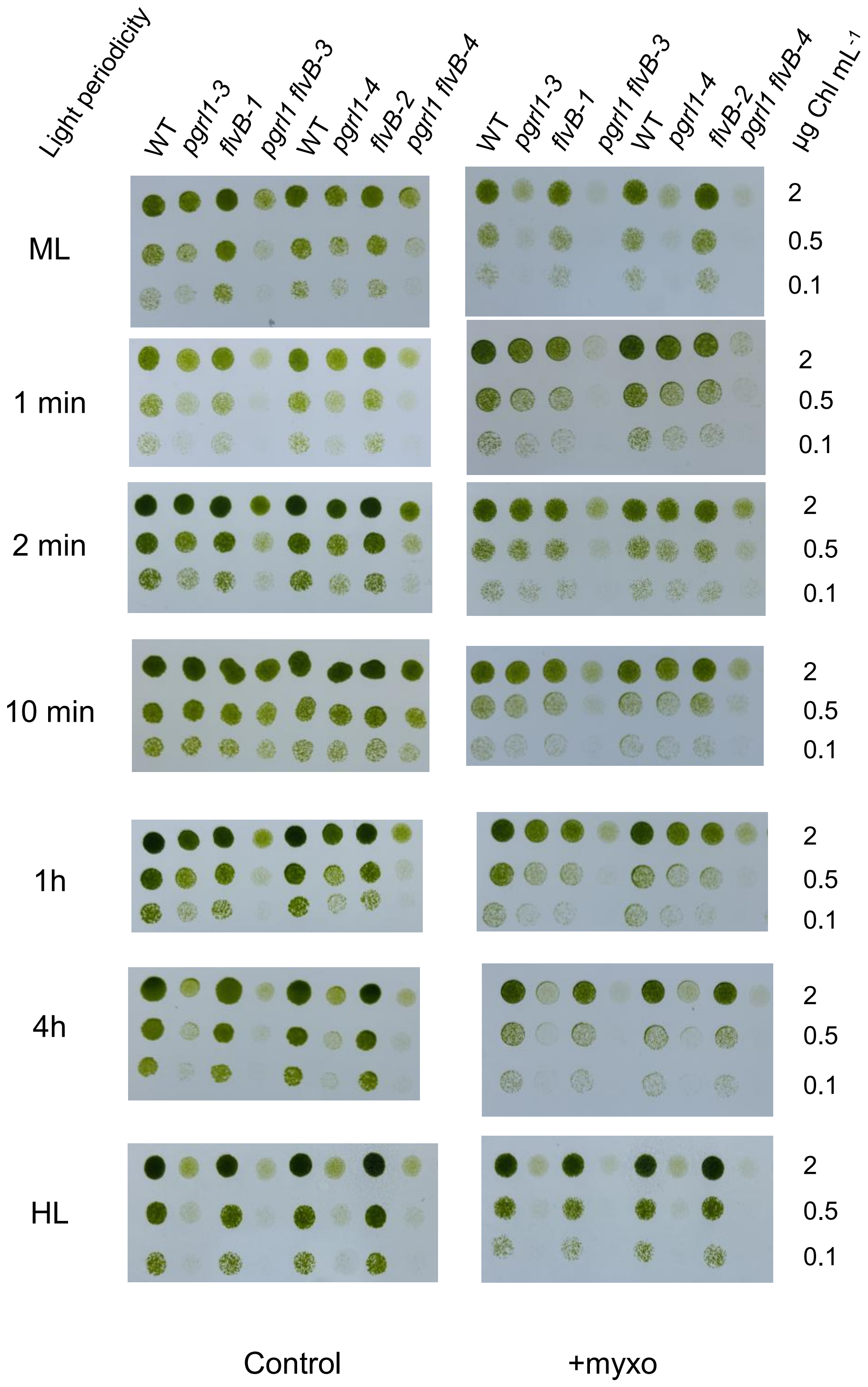


**Supplemental Figure S9. Growth test of *flvB*, *pgrl1,* and double mutants under various periodicities of light fluctuation.** Growth test of *flvB*, *pgrl1,* and *pgrl1 flvB* double mutants (-3 and -4) and their control strain under periodic darkness/high light (0/100 µmol photon $m^{-2}$ $s^{-1}$) shifts with various periodicities or continuous medium (ML, 50 µmol photon $m^{-2}$ $s^{-1}$) or high (HL, 100 µmol photon $m^{-2}$ $s^{-1}$) light. Cells were spotted on plates containing minimal medium at pH 7.2 and grown under the various illumination patterns in the absence (left panels) or presence (right panels) of 2.5 $\mu$M myxothyazol in air enriched at 2.5 % $CO_2$. Cells were left to grow between 5 to 11 days (**Table S.5**). Spots shown are representative of $n$=5 biologically independent experiments. Supports **Fig. 2**.

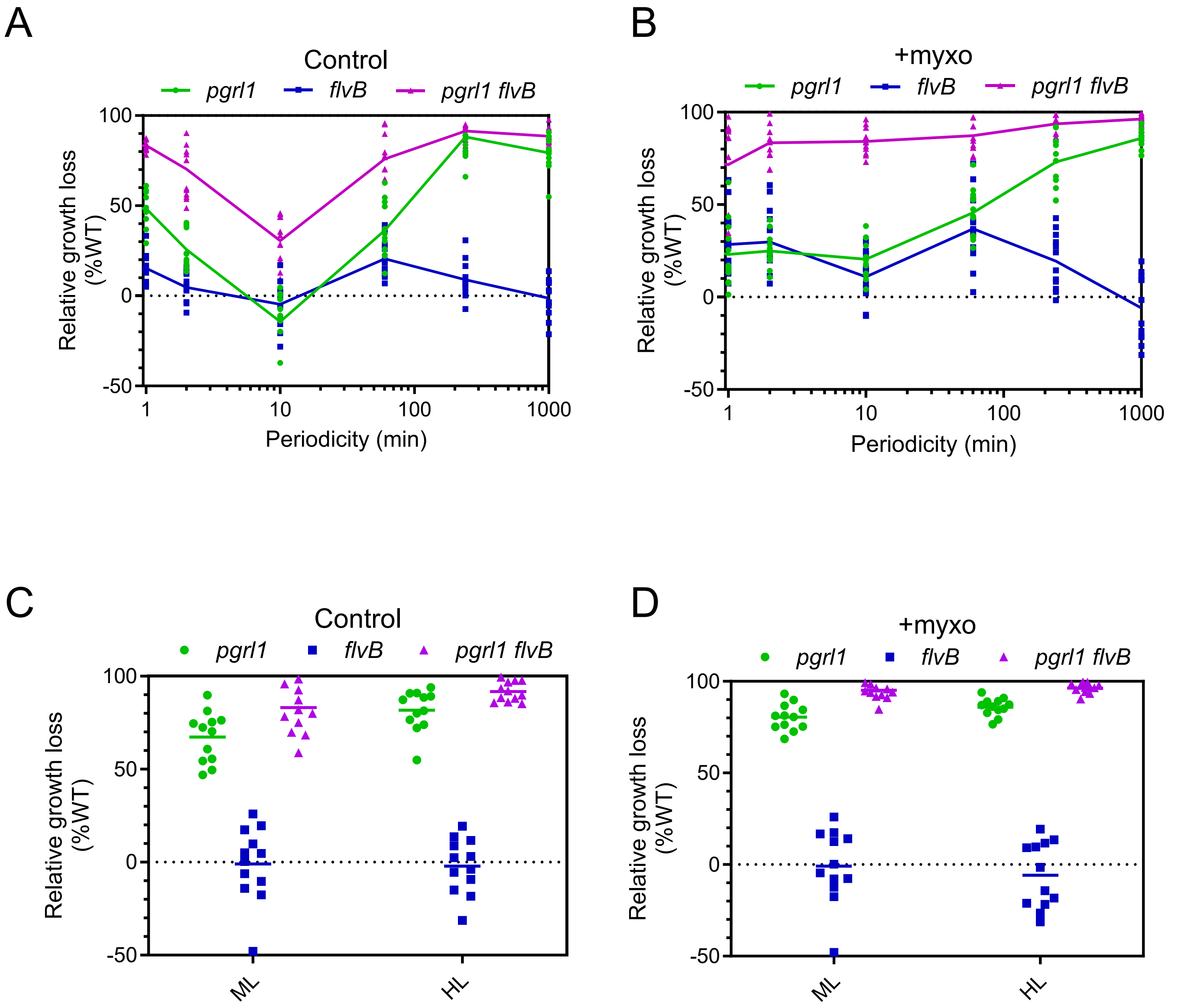


**Supplemental Figure S10. Growth loss linked with *pgrl1*, *flvB*, and *pgrl1 flvB* mutation.** (**A, B, C, D**) Quantification of the loss of growth capacity relative to the WT levels in *pgrl1*, *flvB,* and *pgrl1 flvB* mutants in the absence (**A, C**) or presence (**B, D**) of myxothyazol (2.5 $\mu$M final concentration) and for various periodicities of light (**A, B**) or continuous medium (ML, 50 $\mu$mol photon $m^{-2}$ $s^{-1}$) or high (HL, 100 $\mu$mol photon $m^{-2}$ $s^{-1}$) light (**C, D**). Shown are greenness levels relative to the WT for the first two dilutions of spots shown in **Figs. S5,S6**. Plain lines connect averages, and dots represent individual replicates ($n$ = 2 technical replicates for each $n$ = 6 biologically independent samples). Note that replicates from *pgrl1-2* are not shown here as this strain showed a stronger general sensitivity to light, independent of light periodicity, which was not linked to the *pgrl1* mutation (**Figs. S5-S8**). Show at a periodicity of 1000 minutes is continuous HL. Supports **Fig. 2**.

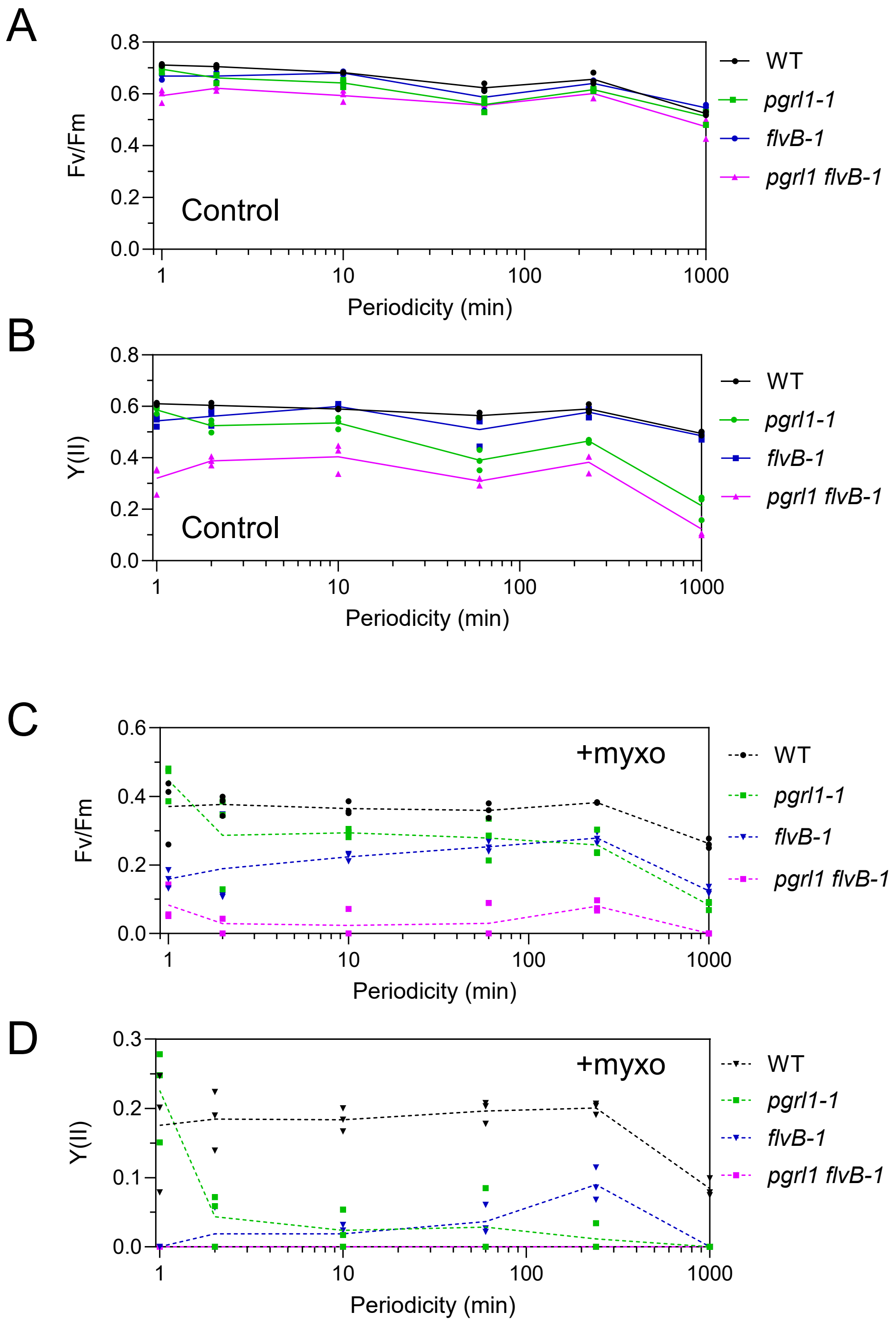


**Supplemental Figure S11. Fv/Fm and Y(II) of *flvB*, *pgrl1,* and *pgrl1 flvB* mutants.** (**A, B, C, D**) Maximal PSII yield (Fv/Fm, **A**, **C**) and operating PSII yield under low light (Y(II), **B**, **D**) were measured after receiving a number of photons equivalent to 4 hours of continuous high light (100 $\mu$mol photon $m^{-2}$ $s^{-1}$) for a range of light periodicities, on *pgrl1-1*, *flvB-1*, *pgrl1 flvB-1* mutants and their control strain (WT) in the absence (**A, B**) or presence (**C, D**) of myxothyazol (2.5 $\mu$M final concentration). **B** and **D** show PSII yield (Y(II)) for the same samples as in **A** and **C,** respectively. Plain lines connect averages, and dots represent individual replicates ($n$ = 3 biologically independent samples). Supports **Fig. 3**.

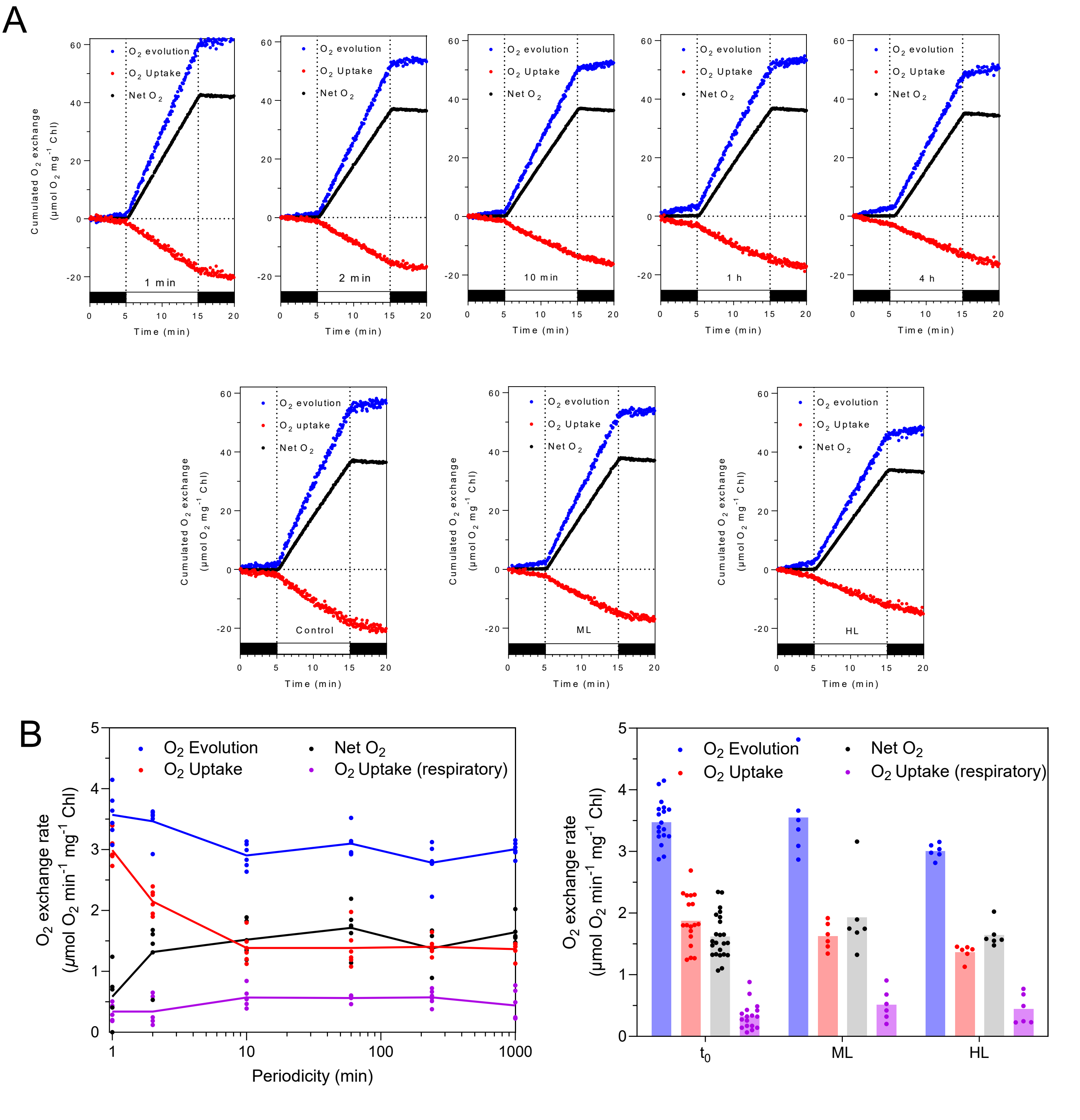


**Supplemental Figure S12. Gas exchange characteristics of WT cells acclimated to various periodicities of light.** Gas exchange was measured in wildtype cells after receiving a number of photons equivalent to 4 hours of continuous high light (100 *µ*mol photon m$^{-2}$ s$^{-1}$) for a range of light periodicities. (**A**) Cumulated $O_2$ exchange upon a dark-light-dark transition on cells acclimated to each light periodicity (1 min- 4 h), medium light (ML, 50 *µ*mol photon m$^{-2}$ s$^{-1}$), high light (HL, 100 *µ*mol photon m$^{-2}$ s$^{-1}$), and before the treatment (Control). Shown are representative cumulative $O_2$ exchange ($n$ = 6 biologically independent samples). (**B**). $O_2$ exchange rates measured during the first minute of a dark to light transition (Gross evolution, Gross uptake, and Net evolution) and initial dark respiration (Uptake) for the range of light periodicities (left panel) or continuous ML, HL, and control conditions ($t_0$). Dots represent individual replicates ($n \geq 6$ biologically independent samples). Supports **Fig. 4**.

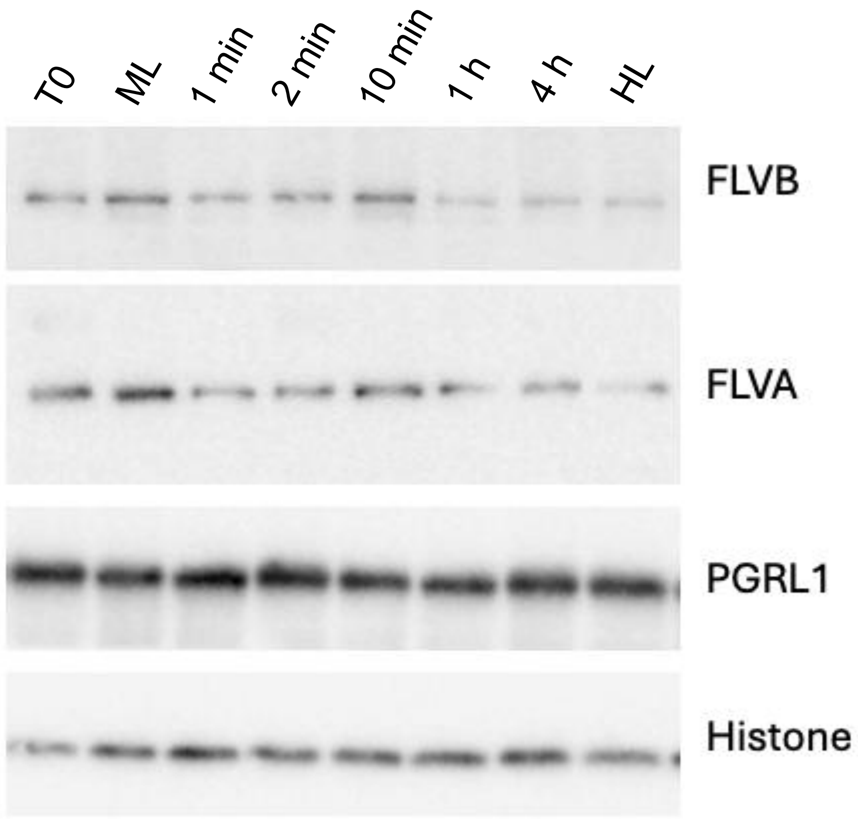


**Supplemental Figure S13. Protein abundance of FLVs and PGRL1 in response to a range of light periodicities.** Immunodetection of FLVA, FLVB, and PGRL1 proteins in wildtype cells (WT), after receiving a number of photons equivalent to 4 hours of continuous high light (HL, 100 $\mu$mol photon m$^{-2}$ s$^{-1}$) for a range of light periodicities and constant HL or medium light (ML, 50 $\mu$mol photon m$^{-2}$ s$^{-1}$). Shown on the left column is the starting level of proteins under our standard growing conditions (T0, 50 $\mu$mol photon m$^{-2}$ s$^{-1}$). Immunodetection of Histone is shown as the loading control. Shown are representative samples ($n$ = 3 biologically independent samples). Samples used are from the same cultures used for **Fig. S12**. Supports **Fig. 4**.

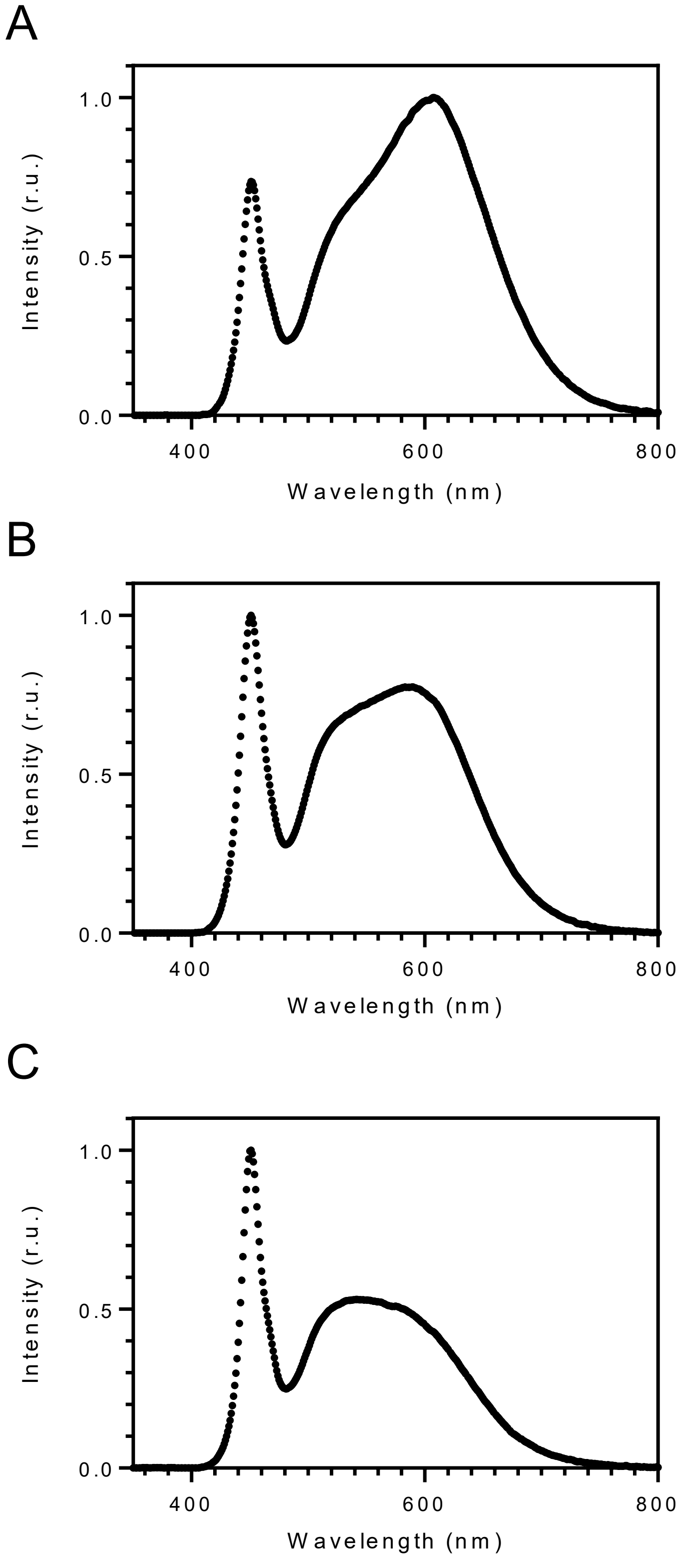


**Supplemental Figure S14. Light spectrum used in this work.** (**A**) Light spectrum of the growth chambers used for preculturing cells for all experiments. (**B**) Light spectrum of the LED panels used during short-term acclimation experiments shown in **Figs. 3, 4, S9-S11**. (**C**) Light spectrum of the LED panels used for solid growth experiments shown in **Figs. 1, 2, S2, S4, S5, S6 and S7**. Supports **Figs. 1-4**.
